# Follicular helper T cell signature of replicative exhaustion, apoptosis and senescence in common variable immunodeficiency

**DOI:** 10.1101/2021.06.15.448353

**Authors:** Giulia Milardi, Biagio Di Lorenzo, Jolanda Gerosa, Federica Barzaghi, Gigliola Di Matteo, Maryam Omrani, Tatiana Jofra, Ivan Merelli, Matteo Barcella, Francesca Ferrua, Francesco Pozzo Giuffrida, Francesca Dionisio, Patrizia Rovere-Querini, Sarah Marktel, Andrea Assanelli, Simona Piemontese, Immacolata Brigida, Matteo Zoccolillo, Emilia Cirillo, Giuliana Giardino, Maria Giovanna Danieli, Fernando Specchia, Lucia Pacillo, Silvia Di Cesare, Carmela Giancotta, Francesca Romano, Alessandro Matarese, Alfredo Antonio Chetta, Matteo Trimarchi, Andrea Laurenzi, Maurizio De Pellegrin, Silvia Darin, Davide Montin, Rosa Maria Dellepiane, Valeria Sordi, Vassilios Lougaris, Angelo Vacca, Raffaella Melzi, Rita Nano, Chiara Azzari, Lucia Bongiovanni, Claudio Pignata, Caterina Cancrini, Alessandro Plebani, Lorenzo Piemonti, Constantinos Petrovas, Maurilio Ponzoni, Alessandro Aiuti, Maria Pia Cicalese, Georgia Fousteri

## Abstract

**Background:** Common variable immunodeficiency (CVID) is the most frequent primary antibody deficiency. A significant number of CVID patients are affected by various manifestations of immune dysregulation such as autoimmunity. Follicular T cells cells are thought to support the development of CVID by providing inappropriate signals to B cells during the germinal center (GC) response.

**Objectives:** We determined the possible role of follicular helper (Tfh) and follicular regulatory T (Tfr) cells in patients with CVID by phenotypic, molecular, and functional studies.

**Methods:** We analyzed the frequency, phenotype, transcriptome, and function of circulating Tfh cells in the peripheral blood of 27 CVID patients (11 pediatric and 16 adult) displaying autoimmunity as additional phenotype and compared them to 106 (39 pediatric and 67 adult) age-matched healthy controls. We applied Whole Exome Sequencing (WES) and Sanger sequencing to identify mutations that could account for the development of CVID and associate with Tfh alterations.

**Results:** A group of CVID patients (*n=9*) showed super-physiological frequency of Tfh1 cells and a prominent expression of PD-1 and ICOS, as well as a Tfh RNA signature consistent with highly active, but exhausted and apoptotic cells. Plasmatic CXCL13 levels were elevated in these patients and positively correlated with Tfh1 cell frequency, PD-1 levels, and an elevated frequency of CD21^lo^CD38^lo^ autoreactive B cells. Monoallelic variants in *RTEL1*, a telomere length- and DNA repair-related gene, were ideintified in four patients belonging to this group. Lymphocytes with highly shortened telomeres, and a Tfh signature enriched in genes involved in telomere elongation and response to DNA damage were seen. Histopathological analysis of the spleen in one patient showed reduced amount and size of the GC that, unexpectedly, contained an increased number of Tfh cells.

**Conclusion:** These data point toward a novel pathogenetic mechanism in a group of patients with CVID, whereby alterations in DNA repair and telomere elongation might be involved in GC B cells, and acquisition of a Th1, highly activated but exhausted and apoptotic phenotype by Tfh cells.

## Introduction

Common Variable Immunodeficiency (CVID) is the most common primary antibody deficiency (PAD) in humans. CVID is characterized by low levels of IgG, IgA, and/or IgM, failure to produce antigen-specific antibodies and accounts for the majority (57%) of symptomatic primary immunodeficiencies according to the European Society for Immunodeficiencies registry (1–3), with estimated prevalence at 1 in 20,000-50,000 new births (www.esid.org). Secondary clinical features of CVID include combinations of various infectious, autoimmune and lymphoproliferative manifestations that complicate the course and the management of the disease. Mortality increases by 11-fold if any secondary clinical feature is present, with an overall survival considerably lower than the general population (4). Ig supply is the mainstay of treatment for CVID that is often combined with immunomodulatory drugs to improve the management of the secondary complications (5). Different autoimmune (AI) manifestations often coincide in the same CVID patient and their management remains a great challenge (6).

Failure of antibody production in CVID can be the direct result of B-cell insufficiency and dysfunction or can be T-cell mediated. For instance, a reduction in the number and percentage of isotype-switched B cells (4–5) as well as a loss of plasma cells in the bone marrow and mucosal tissues has been reported (9); on the other hand, T follicular helper (Tfh) cells, which drive T cell-dependent humoral immunity in germinal centres (GCs), were shown to underpin CVID development in some patients (10). A predominant Th1 phenotype and altered Tfh function have beed described in patients affected by CVID with various manifestations of immune dysregulation (11). Follicular regulatory T cells (Tfr) safeguard the function of Tfh cells limiting AI and excessive GC reactions. The role of Tfr cells in CVID remains unexplored: these cells could be dysfunctional promoting AI or hyperactive over-inhibiting Tfh cells and ultimately leading to CVID.

Next generation sequencing has led to the discovery of an increasing number of monogenic causes of CVID (12,13). In the small list of monogenic CVID disorders, mutations in Tfh-associated genes s such as *ICOS* (14–16), *Il-21* (15,16), SLAM family proteins (17) and others (18) reduce the number or function of Tfh cells. Moreover, activated circulating Tfh cells have been associated with the immune phenotype of patients affected by AI diseases and underpin the production of autoantibodies (AAb) (19–21). On these grounds, we analyzed the frequency, subset distribution, phenotype, transcriptome, and function of Tfh cells in 27 patients with CVID presenting AI as a secondary phenotype. We also investigated Tfr cells and B cells for their frequency and phenotype and determined CXCL13 plasma levels. Finally, we performed WES in a fraction of patients and identified mutations and genetic variants that could account for the development of CVID and their Tfh-related immunophenotype.

## Materials and Methods

### Study cohort

The study encompassed 27 patients that satisfied the following inclusion criteria: increased susceptibility to infection, autoimmune manifestations, granulomatous disease, unexplained polyclonal lymphoproliferation, affected family member with antibody deficiency (low IgA or IgM or IgG or pan-hypogammaglobulinemia), poor antibody response to vaccines (and/or absent isohaemagglutinins), low amount of switched memory B cells, diagnosis established after the 4^th^ year of life, no evidence of profound T-cell deficiency (based on CD4 numbers per microliter: 2-6y <300, 6-12y<250, >12y<200), % naive CD4: 2-6y<25%, 6-16y<20%, >16y<10%), and altered T-cell proliferation (24). CVID was diagnosed according to ESID criteria (25). The study cohort was enrolled in a study conducted at San Raffaele Hospital (HSR) in Milan and was constituted by patients diagnosed at HSR or referred from other centres of the Italian Associazione Italiana Ematologia Oncologia Pediatrica-Associazione Immunodeficienze Primitive (AIEOP-IPINET) (Federico II University of Naples, Ancona University, Bologna University, University of Rome Tor Vergata and Pediatric Hospital Bambin Gesù, Parma Hospital, Meyer Pediatric Hospital in Florence, Alessandria Hospital, University of Brescia, and Regina Margherita Hospital in Turin). The cohort was composed of 16 adults (mean age 36 years, age range 20-63 years) and 11 children (mean age 13 years, age range 6-17 years) (**Table I**). Blood and tissues samples for the study were collected between February 2016 and January 2020, they were compared with 106 age-matched control subjects, 67 of which adults (mean age 27 years, age range 18-52 years) and 39 children (mean age 11 years, age range 2-16 years) (**Table I, E1**). Spleen tissue was collected from the pancreata of non-diabetic brain-dead multiorgan donors received at the Islet Isolation Facility of San Raffaele Hospital, following the recommendation approved by the local ethics committee. Collection of biological specimens was performed after subjects or parents’ signature of informed consent for biological samples collection, including genetic analyses, in the context of protocols approved by the Ethical Committee of HSR (Tiget06, Tiget09 and DRI004 protocols). Tonsils were collected from non-diabetic brain-dead multiorgan donors received at the Islet Isolation Facility of San Raffaele Hospital, following the recommendation approved by the local ethics committee.

**TABLE I.**
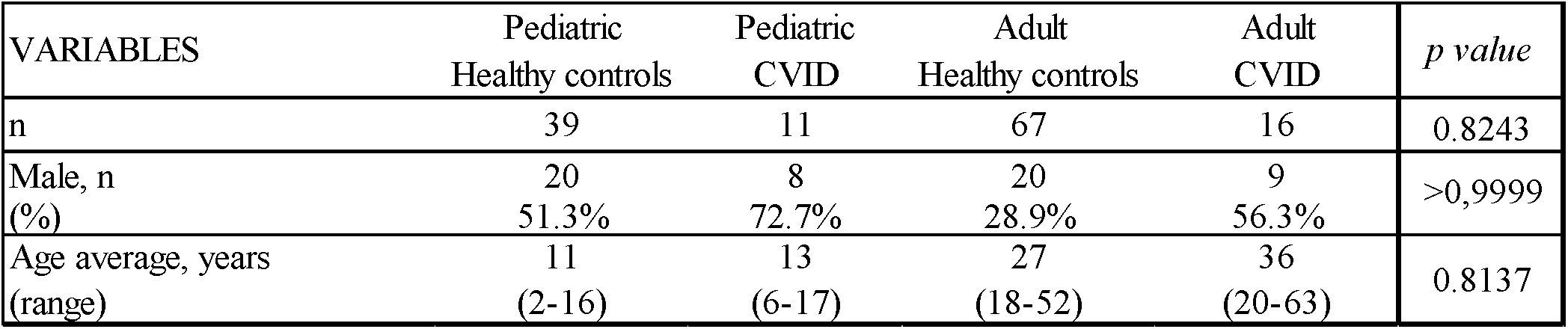
CVID patients and healthy controls

| VARIABLES | Pediatric<br>Healthy controls | Pediatric<br>CVID | Adult<br>Healthy controls | Adult<br>CVID | *p value* |
| --- | --- | --- | --- | --- | --- |
| n | 39 | 11 | 67 | 16 | 0.8243 |
| Male, n<br>(%) | 20<br>51.3% | 8<br>72.7% | 20<br>28.9% | 9<br>56.3% | >0,9999 |
| Age average, years<br>(range) | 11<br>(2-16) | 13<br>(6-17) | 27<br>(18-52) | 36<br>(20-63) | 0.8137 |

### Sample collection, cell staining and flow cytometry

FACS stainings were performed on peripheral blood mononuclear cells (PBMC), whole blood (EDTA), and spleen. PBMC were isolated from heparinized blood by Ficoll density gradient centrifugation using Lymphoprep (Stemcell). Surface staining was performed in fresh isolated PBMC with a panel of mAbs including: CD45RA-PE (HI100, Miltenyi Biotec), CD4-PercP (VIT4, Miltenyi Biotec), CD25-APC (2A3, BD Biosciences), PD-1-PE-Cy7 (eBioJ105, eBiosciences), CD3-APC-Vio770 (BW264/56, Miltenyi Biotec), CXCR5-Brilliant Violet 421 (J252D4, BioLegend), CD19-PO (SJ25C1, BD Biosciences), CD14-VioGreen (TUK4, Miltenyi Biotec), CD8-VioGreen (BW135/80, Miltenyi Biotec) (**Table E2**, staining panel A). Cells were fixed and permeabilized for intracellular staining with the FoxP3/Transcription Factor Staining Buffer Set (eBioscience) prior to staining with FoxP3-Alexa Fluor 488 (259D, BD Biosciences) (**Table E2**, staining panel A).

Surface staining was performed in whole blood (EDTA) with two panels of mAbs. The first panel consisted of: CD45RA-FITC (T6D11, Miltenyi Biotec), CD4-PE (REA623, Miltenyi Biotec), CCR6-PerCP (G034E3, BioLegend), CXCR3-APC (1C6, BD Biosciences), ICOS-PECy7 (ISA-3, Invitrogen), CD3-APC-Vio770 (BW264/56, Miltenyi Biotec), CXCR5-Brilliant Violet 421 (J252D4, BioLegend), CD45-Brilliant Violet 510 (HI30, BioLegend) (**Table E2**, staining panel B). The second panel consisted of: CD45RA-FITC (T6D11, Miltenyi Biotec), PD1-PE (J43, ThermoFisher), CD4-PercP (VIT4, Miltenyi Biotec), CXCR3-APC (1C6, BD Biosciences), ICOS-PECy7 (ISA-3, Invitrogen), CD3-APC-Vio770 (BW264/56, Miltenyi Biotec), CXCR5-Brilliant Violet 421 (J252D4, BioLegend), CD45-Brilliant Violet 510 (HI30, BioLegend) (**Table E2**, staining panel C).

B cell staining was performed on frozen PBMCs after thawing in complete medium (RPMI 10% FBS, PS/G 1X) containing DNase I (Calbiochem, cod 260913) for 10 min at 37°C. 2×10^5^ PBMCs were stained with the following mix of mAbs, as previously described (26,(26)(26)(26)(26)(26)(26)(26)(26)(26)(26)(26)(26)(26)27): CD27-APC (M-T27, BD Biosciences), CD19-PECy7 (SJ 25C1, BD Biosciences), CD21-PE (B-LY4, BD Biosciences), CD38-PerCP5.5 (HIT2, BD Biosciences), CD24-Pacific Blue (SN3, EXBIO), IgD-biotinylated (IA6-2, BD Bioscience), IgM-FITC (G20-127, BD Biosciences), IgG-FITC (polyclonal, Jackson Immunoresearch), IgA-FITC (polyclonal, Jackson Immunoresearch), and Streptavidin-Pacific Orange (ThermoFisher) (**Table E3**, staining panels A-C).

Single cell suspensions from spleen were prepared as previously described (28). B cell and Tfh cell staining was performed on frozen splenocytes as described above (**Table E3**, staining panels A-D). Cells were acquired on FACS CantoII (BD) and analyzed with FlowJo (Tree Star) software.

Single cell suspensions from tonsil were prepared as already described (Efficient Isolation Protocol for B and T Lymphocytes from Human Palatine Tonsils. DOI: 10.3791/53374). Sorting of Centrocytes (CCs), Centroblasts (CBs) and Tfh cells from tonsils (*n* = 3) was performed on frozen cells in complete medium (RPMI 10% FBS, PS/G 1X). Cells were stained with the following mix of mAbs, as previously described (26,27): CD3-APC-Cy7 (BW264/56, Miltenyi Biotec), CD4-PerCP (VIT4, Miltenyi Biotec), CXCR5-BV421 (J252D4, BioLegend), PD1-PE (J43, ThermoFisher), CD27-APC (M-T27, BD Biosciences), CD38-PerCPCy5.5 (HIT2, BD Biosciences), CXCR4-PE-Cy7 (12G5, Miltenyi Biotec) (**Table E3**, staining panels E). Cells were sorted on FACS Aria Fusion (BD) and analyzed with FlowJo (Tree Star) software.

### CXCL13 ELISA assay

CXCL13 was evaluated in plasma EDTA by ELISA (Human CXCL13/BLC/BCA-1 Quantikine® ELISA Kit, R & D Systems) following the manufacturer’s instructions. To collect plasma, whole PB (EDTA) was centrifuged at 1000 rpm for 15 min. Plasma was further centrifuged at 13000 rpm for 10 min to remove debris.

### RNA and DNA extraction

Sorted Tfh cells were resuspended in 200 uL of Trizol (Ambion) and frozen at −80°C. After thawing, 100 uL of chloroform was added and RNA was extracted using the RNeasy Mini Kit (QIAGEN) with a small modification as exemplified here: after an incubation for 2 min at RT, samples were centrifuged at 12000 x g for 15 min at 4°C. The transparent upper phase was transferred to a new tube and an equal volume of 70% ethanol was added. The samples were transferred to a RNeasy Mini spin column and centrifuged for 15 sec at 8000 x g. 350 µL Buffer RW1 was added to the column and spinned down 15 sec at 8000 x g. DNases were inactivated using a mix consisting of 10 µL QIAGEN DNase I with 70 uL Buffer RDD. After an incubation of 30 min at RT with the DNase mix, a second wash was carried out with Buffer RW1. Subsequently, RNA was washed twice with Buffer RPE. Finally, RNA was eluted with 30 µL RNase-free water. DNA was extracted from 200ul whole blood (EDTA) using the QIAmp DNA Blood Mini kit according to the manufacturer’s instructions (Qiagen). RNA and DNA concentration were quantified by NanoDrop 8000 (Thermo Scientific).

### RNA sequencing and analysis

RNA-seq data were trimmed to remove Illumina adapters and low-quality reads using cutadapt. Sequences were then aligned to the human reference genome (GRCh38/hg38) using STAR, with standard input parameters. Gene counts were produced using Subread featureCounts, using Genecode v31 as reference. Transcript counts were processed using edgeR, using standard protocols as reported in the manual. Differential expression was determined considering p-values corrected by FDR including sex as covariate. Heatmaps were produced with pheatmap R package, whereas volcano and scatter plots were produced with ggplot2 R package.

### Immunohistochemistry

Formalin-fixed, paraffin-embedded tissue 3-4 µm thick sections from spleen specimens were stained with haematoxylin-eosin and underwent histopathological assessment. For immunohistochemistry (IHC), the following antibodies were applied: CD20, CD3m, Bcl-2 (**Table E4** for clones and dilutions). IHC was performed using the standard avidin-biotin-peroxidase complex method, as described elsewhere (29). The immunostaining for Bcl-6 (clone GI 191E/A8) was performed with an automated immunostainer (Benchmark XT, Ventana Medical Systems) after heat-induced epitope retrieval, which was carried out using Ventana cell conditioning buffer 1 (CC1) for 60 minutes.

Images were obtained on a Nikkon microscope system with a 40× (NA 1.3) and 20× (NA 0.75) objectives.

### Genomic studies

#### Whole Exome Sequencing (WES)

The Whole Exome Sequencing of genomic DNA was performed by *Genomnia* (http://www.genomnia.com/). DNA libraries were sequenced on a Hiseq 4000 (Illumina) for paired-end bp reads. Sequencing reads were mapped to the reference human genome (UCSC hg19 and hg38) with the Torrent Suite (5.10.0). The bam files generated from two chips were merged with the Combine Alignments utility of the Torrent Suite. The samples were analyzed with the workflow Ion Report Ampliseq Exome Single Sample (germline) version 5.6. The quality of the sequencing was verified with the fastqc software v.0.10.1 e samstat v.1.08.

##### Candidate variants responsible for the disease

The variants noted by Ion Reporter were analyzed highlighting those that fulfilled the following criteria:

- Quality > 40 (to exclude false positives)
- Minor allele frequency MAF < 1% (“rare variants”), or < 5% (“uncommon variants”)
- Variants of candidate genes
- Non-synonymous exonic variants, with a strong impact on the protein sequence, i.e.: indels causing frameshift; variants introducing or eliminating stop codons; missense variants predicted by SIFT and/or PolyPhen as potentially deleterious for the structure and functionality of the protein; variations on splicing sites.

Candidate variants were screened based on the phenotypes and any known inheritance pattern of the patients. When the cases were sporadic without a familial inheritance tendency, we firstly hypothesized that the patients had a monogenic disorder with an autosomal recessive pattern caused by a homozygous or compound heterozygous inheritance or with a de novo or heterozygous dominant mutation with an incomplete penetrance. If no causative mutations were found, we considered the case as a complex form of CVID. Afterwards, top likely disease-associated variants were validated by Sanger sequencing. **Table E5** includes the gene pipeline used for the discovery of possible causative mutations of CVID designed based on known genes or candidate published in the literature, IUIS classification, as previously shown(30).

### Oligonucleotides for PCR and Sanger sequencing

Primers used for amplification and sequencing of genomic DNA are shown in **Table E6**. Amplified DNA fragments were purified using QIAquick PCR Purification Kit (QIAGEN), according to the manufacturer’s instructions. At the end of the purification 400 ng of DNA were sent to sequence with the Sanger method to Eurofins Scientific.

### Sanger sequencing analysis

The electropherogram of each sample obtained from the Sanger sequencing was analyzed using FinchTV program and Nucleotide BLAST, a search nucleotide databases collection.

### Telomere length

Telomere testing was performed by Repeat Diagnostics Inc.

### RTEL1 gene expression assay

RNA was extracted from 2*10^5 sorted CC, CB and Tfh cells (RNeasy Micro Kit, Qiagen), quantified (NanoDrop™ 2000, ThermoFisher) and retrotranscribed (High-Capacity cDNA Reverse Transcription Kit, ThermoFisher). Gene expression was performed in duplicates using a Droplet Digital PCR system (Bio-Rad) following manufacturer’s instructions (BioRad Droplet Digital PCR Applications Guide, Bulletin_6407). The following ddPCR Gene Expression Assay were used in duplex: AL353715.1 – RTEL1, Human, FAM (dHsaCPE5191681); XBP1, Human, HEX (dHsaCPE5033517); BCL6, Human, HEX (dHsaCPE5034897); HPRT1, Human, FAM (dHsaCPE5192871). Data were analysed with QuantaSoft 1.7.4.0917 (Bio-Rad) software.

### Statistics

Statistical analyses were performed using GraphPad Prism software version 7. Quantitative data are expressed as median (range), and categorical data expressed as percentage (percentage). Comparisons between 2 groups were performed using non□parametric Mann–Whitney *U*□tests. Comparisons among > 2 groups were performed using ANOVA test. Relationships between different parameters were examined using Pearson correlation coefficient. Statistical significance of clinical data was assessed with the Fisher exact test. *P* values ≤ 0.05 were considered significant and indicated with an asterisk. **, *** and **** stand for *P* values ≤ 0.01, ≤ 0.001 and ≤ 0.0001, respectively.

## Results

### CVID patients show prevalence of Th1 follicular helper T cells, which induce IgM but no IgG production in *vitro*

We studied the frequency, activation status, and subset distribution of circulating Tfh and Tfr cells in peripheral blood of a cohort of 27 patients (16 adults and 11 pediatrics) with CVID (**Table I**). None of the patients had a genetic diagnosis at the time of recruitment. Patients were compared to 106 age-and sex-matched healthy controls (HC) (**Table I**). Tfh (CXCR5^+^FoxP3^-^), Tfr (CXCR5^+^FoxP3^+^) and Treg (CXCR5^-^FoxP3^+^) cell frequencies were determined on isolated PBMC (Fig 1*A* and Fig E1, *A*, gating strategy in this article’s Online Repository at www.jacionline.org). CXCR5^+^FoxP3^-^ CD4 T cells expressed low or intermediate CD45RA levels suggesting they were antigen experienced (Fig E1, *B*). The frequency of Tfh cells observed in patients was higher than in HC but quite heterogenous (median 13.90% in CVID vs. 10.90% in HC; *p* = 0.0132) (Fig 1*B*). Three patients had a Tfh cell frequency below the lower cut-off seen in HC (3.84%), while four patients had a Tfh cell frequency above the higher cut-off (23.30%) (Fig 1*B*). The remaining patients had a Tfh cell frequency similar to the one observed in HCs (Fig 1*B*). No significant difference was observed in the frequency of Tfr cells (median 1.48% in CVID vs. 1.47% in HC; *p* = 0.5407) (Fig 1*C*). Accordingly, blood Tfh : Tfr cell ratio was higher in CVID patients in comparison to HC (median 9.56 in CVID vs. 7.41 in HC; *p* = 0.0449) (Fig 1*D*).

**FIGURE 1.**
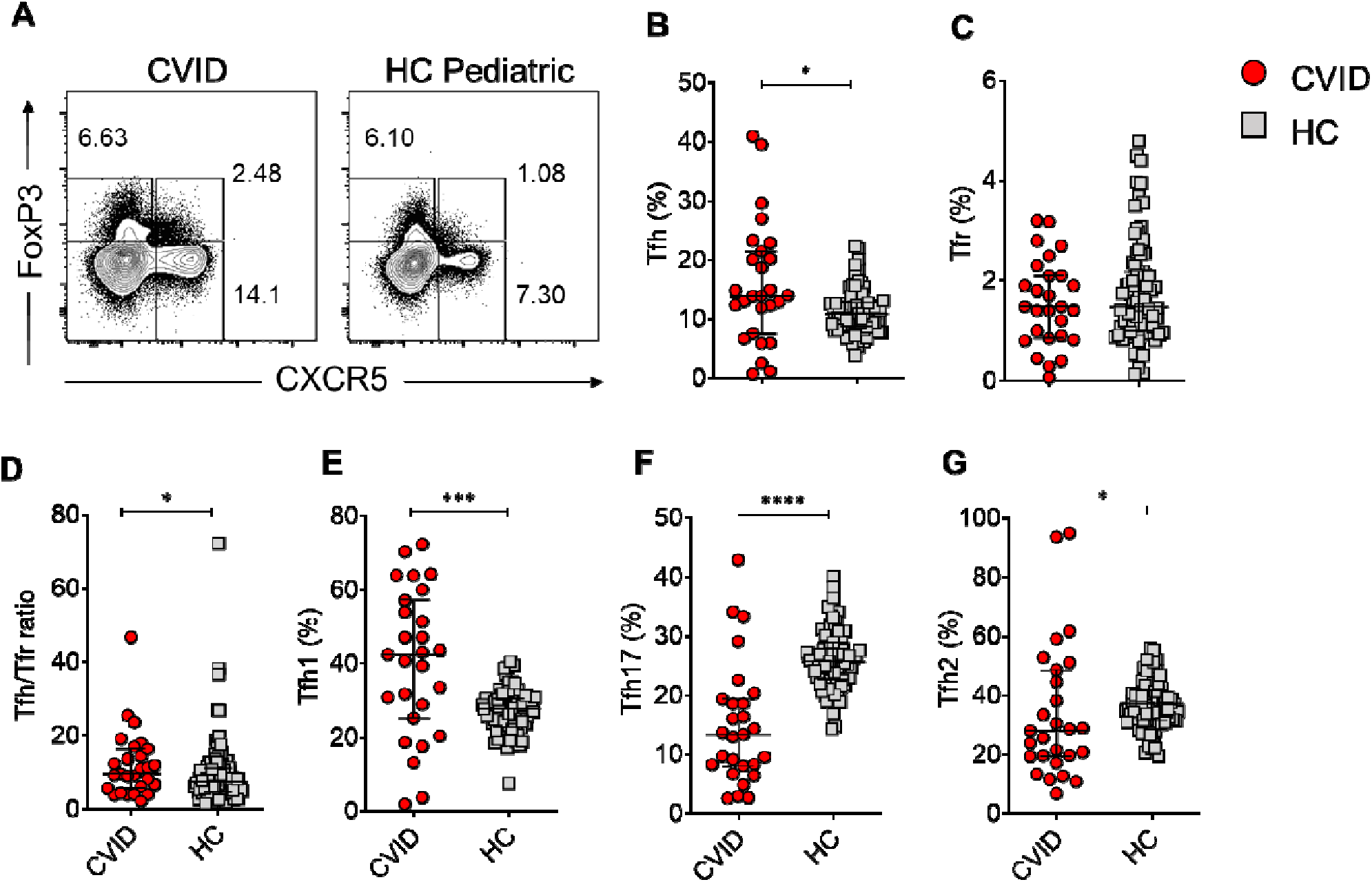
Great variability in percentages of circulating follicular helper T (Tfh) and their subsets in peripheral blood samples from common variable immunodeficiency and autoimmunity (CVID) patients respect to controls. **(A)** Representative flow cytometry plots for Tfh (CXCR5^+^CD4^+^), follicular Treg (FoxP3^+^CXCR5^-^) and conventional Treg (CXCR5^-^CD4^+^), gated on singlets lymphocytes, CD3^+^CD14^-^ CD8^-^CD19^-^. Percentages of Tfh **(B)**, Tfr **(C)**, Tfh:Tfr ratio **(D)** and Tfh subsets in peripheral blood of CVID patients compared to age-matched healthy controls (HC). From left to right: **(E)** frequencies of Tfh1 subset (CXCR3^+^ CCR6^-^), **(F)** Tfh17 (CXCR3^-^ CCR6^+^), **(G)** Tfh2 (CXCR3^-^ CCR6^-^). In all graphs, points represent individual donors and asterisks indicate statistical significance as calculated by Mann Whitney test. Black bars: median with interquantile range. *p<0,05; **p<0,005; ***p<0,001; ****p<0,0001.

Three human Tfh subsets cells can be defined according to the differential expression of CXCR3 and CCR6: CXCR3^+^CCR6^-^ Tfh1 cells, CXCR3^-^CCR6^-^, Tfh2 cells, and CXCR3^-^ CCR6^+^ Tfh17 cells (Fig E1, *C*) (31). Our cohort of CVID patients had a significantly higher percentage of Tfh1 cells as compared to HC (median 42.50% in CVID vs. 27.705 in HC; *p* = 0.0002) (Fig 1*E*), in line with previous reports (12)(33). The percentage of Tfh17 cells was lower in patients as compared to HC (median 13.40% in CVID vs. 25.70% in HC; *p* < 0.0001) (Fig 1*F*), but the proportion of Tfh2 cells was not significantly different between CVID and HC (median 28.10% in CVID vs. 37.20% in HC; *p* = 0.0155) (Fig 1*G*).

We also assessed the frequency of peripheral blood B cell subsets defined by the expression of CD21, CD38, CD24, CD27 and Ig markers in 8 CVID patients and 90 age-and sex-matched HC (Fig *E2A-B* for gating strategy). The frequency of circulating CD19^+^ B cells was reduced in the tested patients as compared to controls (median 1.48% in CVID vs. 10.10% in HC; *p* < 0.0001) (Fig *E2C*). Memory B cells (CD19^+^CD27^+^) were reduced in patients as compared to HC (median 7.89 in CVID vs. 17 in HC; *p* = 0.0062) (Fig *E2D-E*), while no differences were observed in CD19^+^CD27^-^ naive B cells (median 89.40% in CVID vs. 82.10% in HC; *p* = 0.0507), The percentage of CD38^hi^CD24^hi^ transitional B cells in patients was similar to HC (median 4.74% in CVID vs. 7.64% in HC; *p* = 0.3265) (Fig *E2F*). While CVID patients showed no significant difference in the percentage of SM B cells (median 4.35% in CVID vs. 8.18% in HC; *p* = 0.1129) (Fig *E2G*), a lower percentage of CD27^+^IgG^+^ B cells was seen in some patients (median 7.24% in CVID vs. 12.50% in HC; *p* = 0.0709) (Fig *E2H*). Strikingly, a very small percentage of CD27^+^IgA^+^ cells was detected in patients (0% in CVID vs. 10.20% in HC; *p* = 0.0056) (Fig *E2I*). Additionally, CVID patients showed a lower percentage of memory B cells as compared to HC (median 2.54% in CVID vs. 12.92% in HC; *p* <0.0001) (Fig *E2J*), a percentage of autoreactive CD21^lo^CD38^lo^ B cells close to HC (median 1.59% in CVID vs. 2.15% in HC; *p* =0.7700) (Fig *E2K*), and an elevated frequecy of plasma cells (median 10.20% in CVID vs. 0.53% in HC; *p* < 0.0001) (Fig *E2L*).

To assess the functionality of CVID-derived Tfh cells in providing B-cell help *in vitro*, we co-cultured sorted CXCR5^+^CD25^-^ CD4^+^ Tfh cells with naive (CD27^+^ CD38^-^ CD19^+^, B_N_) or memory (CD27^+^ CD38^-^ CD19^+^, B_M_) B cells. B cell differentiation into plasmablasts, IgM and IgG production were evaluated on day 7 of culture. Overall, Tfh cells from five CVID patients showed reduced ability to induce plasmablast differentiation of B_N_ cells (median 54.85% in CVID B_M_+Tfh vs. 70.60% in HC B_M_+Tfh; *p* =0.2621; median 23.40% in CVID B_N_+Tfh vs. 64.40% in HC B_N_+Tfh; *p* =0.0987) (Fig *E2,M*). IgG was almost undetectable in all CVID Tfh : B cell co-cultures irrespective the type of B cells (median 63.18 ng/mL in CVID B_M_+Tfh vs. 9385 ng/mL in HC B_M_+Tfh; p < 0.0001; median 0 ng/mL in CVID B_N_+Tfh vs. 4087 ng/mL in HC B_N_+Tfh; p =0.0029) (Fig *E2,N*). However, both B_N_ and B_M_ cells from CVID patients were able to secrete IgM when cocultured with autologous Tfh cells (median 7781 ng/mL in CVID B_M_+Tfh vs. 1299 ng/mL in HC B_M_+Tfh; *p* = 0.0007; median 4996 ng/mL in CVID B_N_+Tfh vs. 1149 ng/mL in HC B_N_+Tfh; *p* = 0.3922) (Fig *E2,O*). Taken together, a Th1 Tfh cellular shift, reduced frequency of memory B cells, and an inability to class-switch were observed in our cohort of patients with CVID.

### Predominance of Tfh1^hi^Tfh17^lo^PD-1^hi^CXCL13^hi^ immunophenotype in a group of CVID patients

PD-1 and ICOS are the main markers of Tfh cell activation and are considered indicators of their functional status (34). Here, we analyzed the frequency of PD-1- and ICOS-expressing Tfh cells, and the proportion of the highly functional (HF) Tfh subset (CXCR3^-^PD-1^+^). Overall, Tfh cells from CVID patients showed an increased expression of PD-1 (median 40.50 in CVID vs. 21.55 in HC, *p* < 0.0001) and ICOS (median 2.60 in CVID vs. 1.35 in HC, *p* = 0.0009) as compared to controls (Fig 2*A-C*). A significant fraction of PD-1^+^ Tfh cells was also CXCR3^-^ and, consequently, the percentage of HF Tfh cells was higher in patients when compared to HC (median 11.80 in CVID vs. 10.15 in HC; *p* = 0.0367) (Fig 2*D*). The elevated expression of PD-1 was not restricted to the CXCR3^-^Tfh population as CXCR3^+^ Tfh, Tfr and Treg cells also showed elevated PD-1 expression (data not included).

**FIGURE 2.**
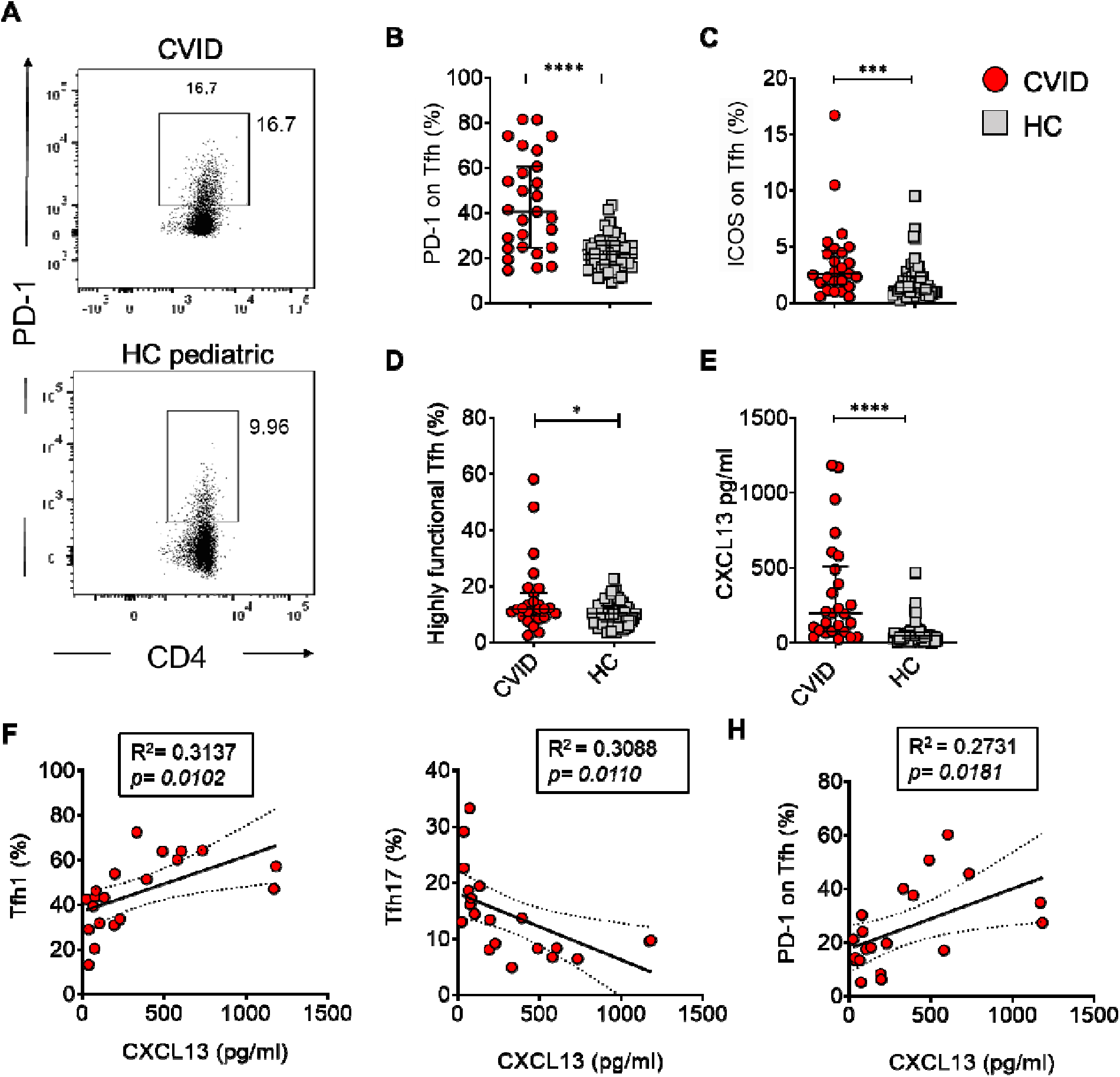
Programmed death (PD)-1 and Inducible co-stimulator (ICOS) expression on circulating Tfh cells is higher in CVID patients compared to controls. Same donors as in Fig. 1 were analyzed. **(A)** Representative flow cytometry plots show PD-1 frequency on CVID patients and pediatric healthy control. **(B-C)** Percentages of PD-1 and ICOS on total Tfh. **(D)** Frequencies of Highly Functional Tfh (CXCR3^-^ PD-1^+^ CXCR5^+^ CD4^+^) cells. **(E)** CXCL13 levels (pg/mL) measured by ELISA assa in plasma of CVID patients compared to controls. Points represent individual donors and asterisks indicate statistical significance as calculated by Mann Whitney test. Black bars: median with interquantile range. *p<0,05; **p<0,005; ***p<0,001; ****p<0,0001. **(F-H)** Correlation analysis between CXCL13 plasma levels and frequencies of Tfh1, Tfh17, PD-1^+^ on Tfh in CVID patients. Frequencies were analyzed by flow cytometry. Lines represent linear regression and SD. *p<0,05; **p<0,005; ***p<0,001; ****p<0,0001.

CVID patients were also characterized by an elevated plasma CXCL13 level as compared to HC (median 196.1 pg/mL in CVID vs. 47.68 pg/mL in HC; *p* < 0.0001) (Fig 2*E*). Furthermore, a significant correlation between CXCL13 plasma levels and the percentage of Tfh1 (Fig 2*F*), and Tfh17 (Fig 2*G*) subsets was seen in patients. In addition, CXCL13 levels correlated positively with the levels of PD-1 expressed by CVID Tfh cells (Fig 2*H*). Interestingly, the percentage of Tfh1 subset showed a negative correlation with the frequency CD19^+^ B cells, while it correlated positively with that of CD21^lo^CD38^lo^ B cells (Fig *E3,A-B*). CD21^lo^CD38^lo^ B cells also showed a positive correlation with plasma CXCL13 levels (Fig *E3,C*).

Based on these findings, we thought to distinguish CVID patients in two groups. We used two criteria: the frequency of Tfh1 cells and used as cut-off a >40% value, the higher value observed in our HC cohort (Fig 3*A*). Another discriminator were the levels of CXCL13 in the plasma (>300pg/ml) (Fig 3*B*). A total of *n=9* patients fulfilled both criteria (Group A). As expected, the frequency of Tfh17 in this group was in the lower range, below the lower cut-off seen in the HC (Fig 3*C*). The majority of Tfh cells of this group also expressed high levels of PD-1 (Fig *D*). The rest of the patients made part of Group B (*n=18*). Hence, based on the frequency of Tfh1 cells and plasma CXCL13 levels, we identified two groups of CVID patients, group A with a prevalence of Tfh1^hi^Tfh17^lo^PD-1^hi^CXCL13^hi^ immunophenotype and group B, with a Tfh-related immunophenotype that was more similar to HC (Fig 3*E*).

**FIGURE 3.**
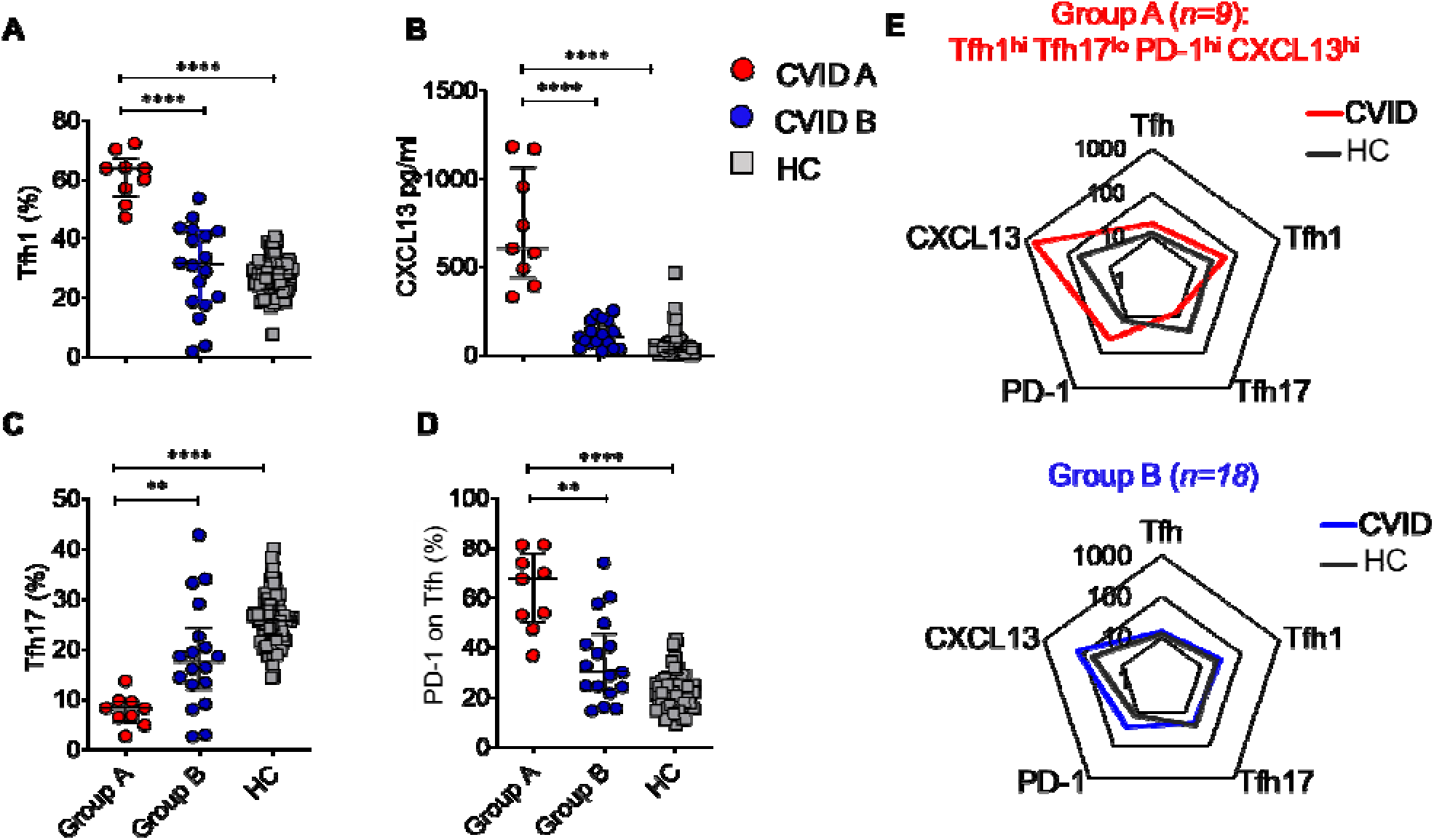
Two major categories of CVID patients based on Tfh-related markers. **(A-D)** Percentages of Tfh1, CXCL13, Tfh17 and PD-1 divides CVID patients in two groups: group A Tfh1^hi^Tfh17^lo^PD-1^hi^CXCL13^hi^ vs group B Tfh1/Tfh17/PD-1/CXCL13^normal^. Percentages were analyzed by flow cytometry. In all graphs, red points represent individual donors of group A and blue points individual donors of group B. Asterisks indicate statistical significance as calculated by Mann Whitney test. Black bars: median with interquantile range. *p<0,05; **p<0,005; ***p<0,001; ****p<0,0001. **(E)** Radar charts represent the percentage of Tfh, Tfh1, Tfh17, PD-1 and CXCL13 in CVID group A vs CVID group B.

Next, we assessed the cytokine and chemokine plasma profile in some CVID patients of group A (*n*=5) and group B (*n*=3) (Fig *E4*). A stronger Tfh1 cell signature (IFN-γ, IP-10, IL-1β, and IL-18) was observed in the majority of group A CVID patients when compared to group B (Fig *E4,A*). Group A patients were also characterized by significantly higher plasma concentrations of chemoattractant proteins CXCL11, CXCL9, and IL-16 (Fig *E4,B*). BAFF, APRIL, CD30, CD40L (Fig *E4,C*), IL-2R (CD25), GCSF, and inflammatory proteins MDC and MIP3a (Fig *E4,D*) were also higher in the plasma of some CVID group A patients. Other cytokine and chemokines were similar between the two groups (Fig *E4, E*).

Clinically, all patients in group A displayed splenomegaly and lymphadenopathy as compared to group B (**Table II**). The prevalence of autoimmune cytopenia, granulomatus disease or enteropathy did not differ between the groups. Hence, we have identified a group of patients characterized by a Tfh1^hi^Tfh17^lo^PD-1^hi^CXCL13^hi^ predominant immunophenotype that had splenomegaly and lymphadenopathy as a common clinical feature.

**TABLE II.** Clinical Features

|  | Group A (*n*=9) | Group B (*n*=18) | *p value* |
| --- | --- | --- | --- |
| **Splenomegaly/lymphadenopathy** | 9 (100%) | 11 (61.1%) | 0,0593 |
| **Autoimmune cytopenia** | 7 (77.77%) | 9 (50%) | 0,2311 |
| **Granuloma** | 2 (22.22%) | 5 (27.77%) | 1 |
| **Pulmonary disease** | 6 (66.66%) | 9 (50%) | 0,4348 |
| **GI disease** | 3 (33.33%) | 3 (16.66%) | 0,3673 |
| **>1 autoimmune diseases** | 3 (33.33%) | 2 (11.11%) | 0,295 |

### Hyperactivated Tfh transcriptional signature characterized by replicative exhaustion and apoptosis in CVID patients with super physiological Tfh1 and CXCL13 levels

Next, we explored the overall transcriptomic landscape of sorted CD4^+^CXCR5^+^CD25^-^ Tfh cells in CVID group A (high Tfh1, CXCL13 high) vs. group B (normal Tfh1 and CXCL13) patients through RNA-Seq. We performed several differential gene expression analyses including group A vs B, group A vs HC, group A vs (B+HC) and group B vs HC in order to better figure out expression modulation across all groups. Comparing CD4^+^CXCR5^+^CD25^-^ Tfh cells of CVID group A with HC we identified 427 differentially expressed genes (DEGs), while comparison to CVID group B 58 DEGs (52 DEGS in common) were seen, leading to the conclusion that group A has a different Tfh expression profile compared to the other two groups (Fig 4*A* and Fig *E4*). Hierarchical group analysis for Tfh-related genes showed that Tfh cells in group A CVID patients were characterized by an increased expression of Tfh-highly active signature (e.g. IL-21, Bcl-6), while group B CVID Tfh cells expressed Tfh-related genes at levels that were comparable to HC (Fig 4*B*). Importantly, increased levels of CXCR5, CXCL13 and genes involved in Tfh lineage specification (as Bcl-6, IL-21, Tox2, ICOS, PD-1) had strong influence in separating the two groups (Fig 4*B*). Furthermore, additional hierarchical analysis that took into consideration T cell activation, exhaustion and cell death pathways revealed an increased representation of these pathways in Tfh cells isolated from patients belonging to group A (Fig 4*C-E*). Thus, *in silico* RNA-Seq analyses confirmed the flow cytometry data and identified a group of CVID patients characterized by highly activated Tfh cell immunophenotype. Tfh cells in this CVID group A patients also expressed a strong T cell activation program but also evidence of cellular exhaustion and apoptosis.

**FIGURE 4.**
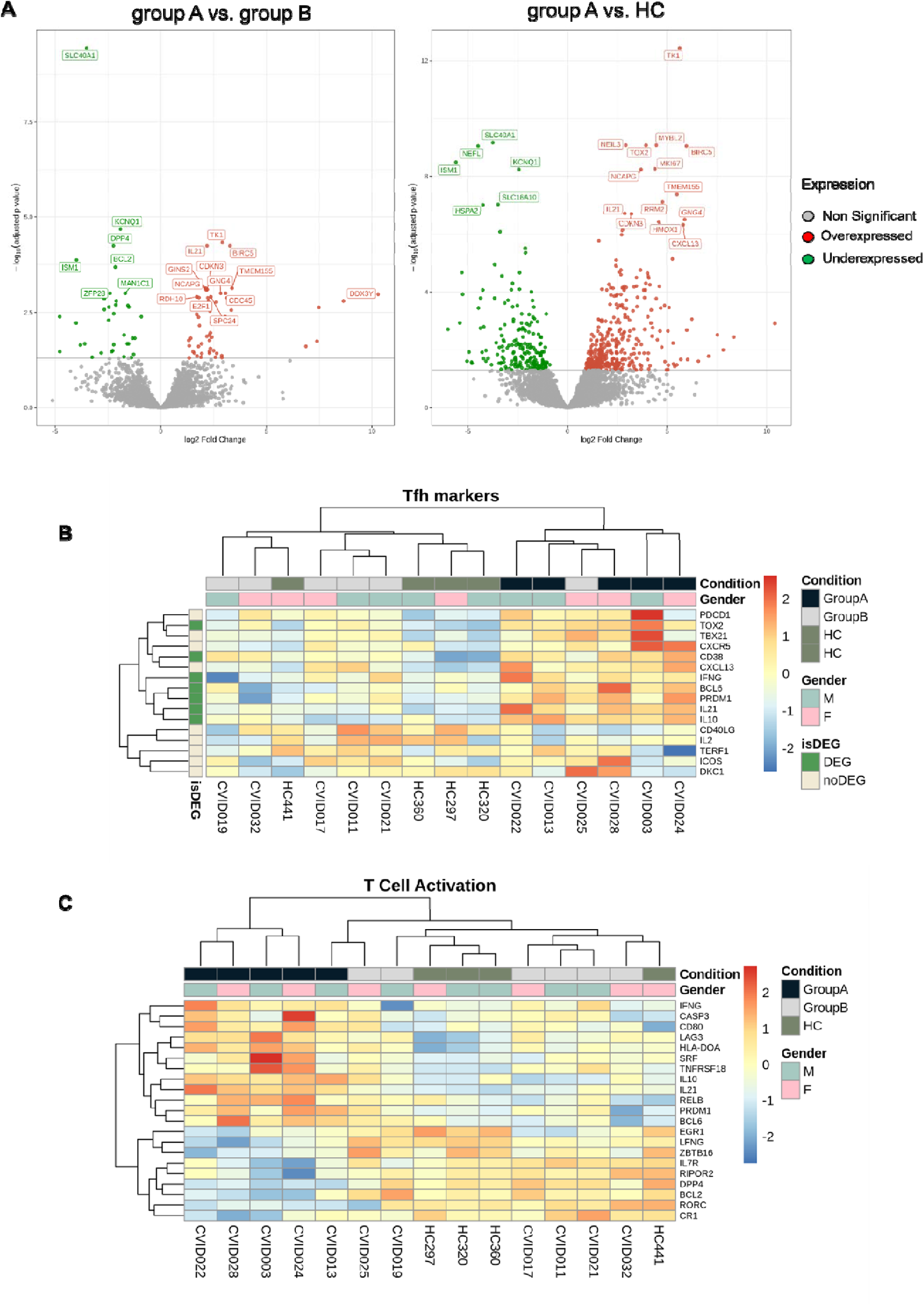
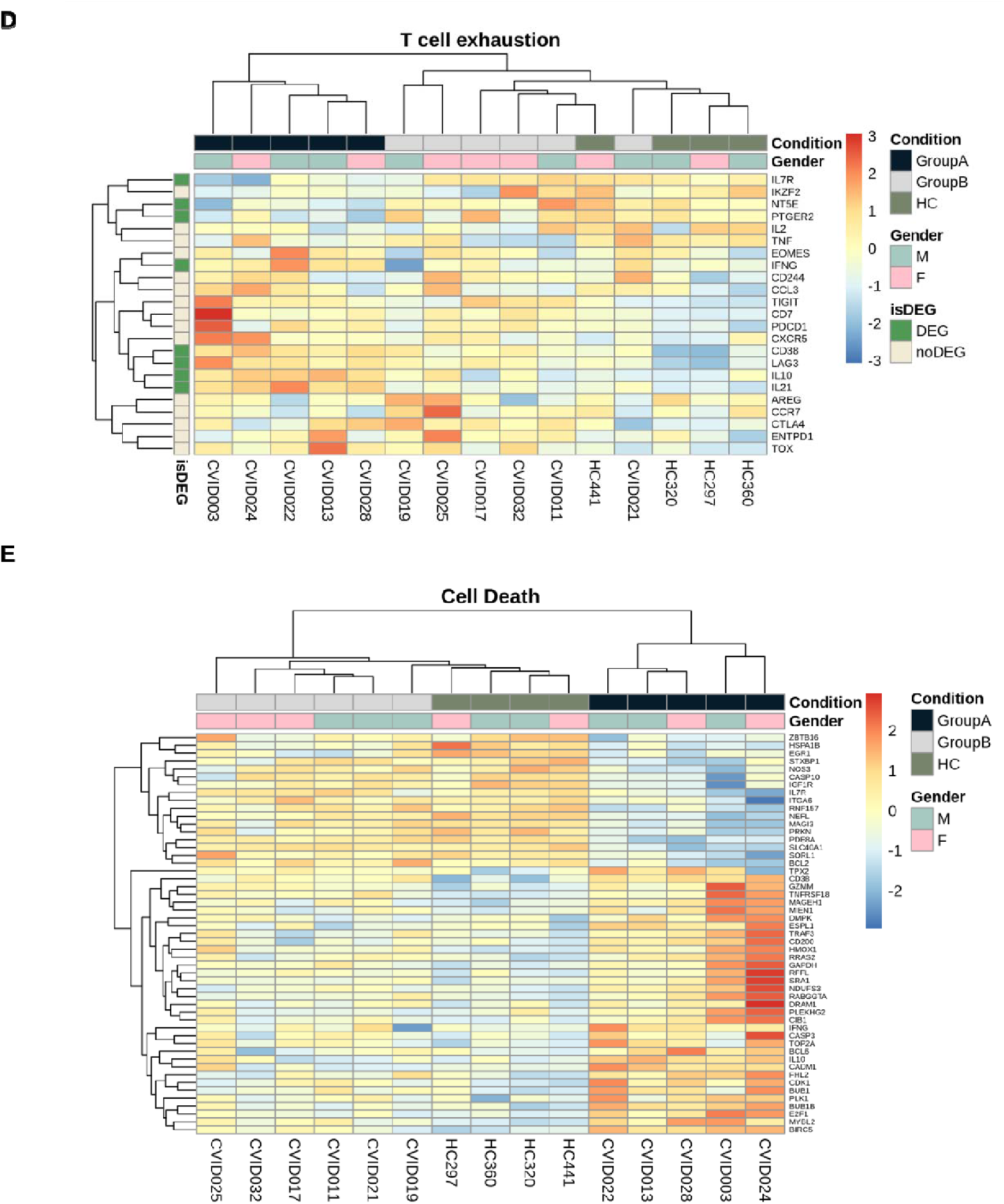
Transcriptomic landscape of sorted CD4^+^CXCR5^+^CD25^-^ Tfh cells in CVID group A vs. group B patients and healthy control carried out through RNA-Seq. **(A)** Vulcano plots representing the gene expression profile of group A vs Group B and group A vs HC. **(B)** Hierarchical grouping analysis based on Tfh-related genes divides patients into Tfh-highly active (group A) and normal (group B). Hierarchical grouping analysis representing the expression of genes involved in **(C)** T cell activation, **(D)** T cell exhaustion and **(E)** cell death pathways. They divide patients into group A and group B. In the volcano plots and heat maps, red color intensities represent a higher gene expression.

### Heterozygous variants in *RTEL1* identified in group A CVID patients associate with senescent lymphocytes

WES analysis of genomic DNA in nine CVID patients, 6 from group A and 3 from group B identified 24 variants in the probands. Patients were prioritized on the basis of the severity of their clinical profile and Tfh1^hi^Tfh17^lo^PD-1^hi^CXCL13^hi^ immunophenotype. Variants were filtered for association with different forms of primary immunodeficiency including CVID (**Table E5**)(30). A total of 16 variants were confirmed by Sanger sequencing in six patients (**Table III**). Of these, heterozygous missense mutations in *RTEL1* gene (c.2785G>A, p.Ala929Thr; c.2123G>A, p.Arg708Gln; c.2051G>A, p. Arg684Gln; c.371A>G, p. Asn124Ser), which is essential for DNA replication, genome stability, DNA repair and telomere maintenance (35–37), were observed in four out of 5 tested CVID patients belonging to group A (**Table III**). Group A patients with *RTEL-1* variants had heterozygous variants in additional genes, i.e., in *PRF1* (c.273G>A, p.Ala91Val), *PRKDC* (c.9503C>T, p.Gly3149Asp), *STXBP2* (c.1331C>T, p.Ala444Val), *MST1* (c.1012T>C, p.Cys338Arg), *TNFRSF13B* (c.659T>C, p.Val220Ala), and *LYST* (c.2433C>T, p.Ser753Asn), suggesting that their disease could be influenced by other heterozygous mutations in association with *RTEL1* variant. The immunological and clinical features of each patient are included in **Table E7**.

**TABLE III.**
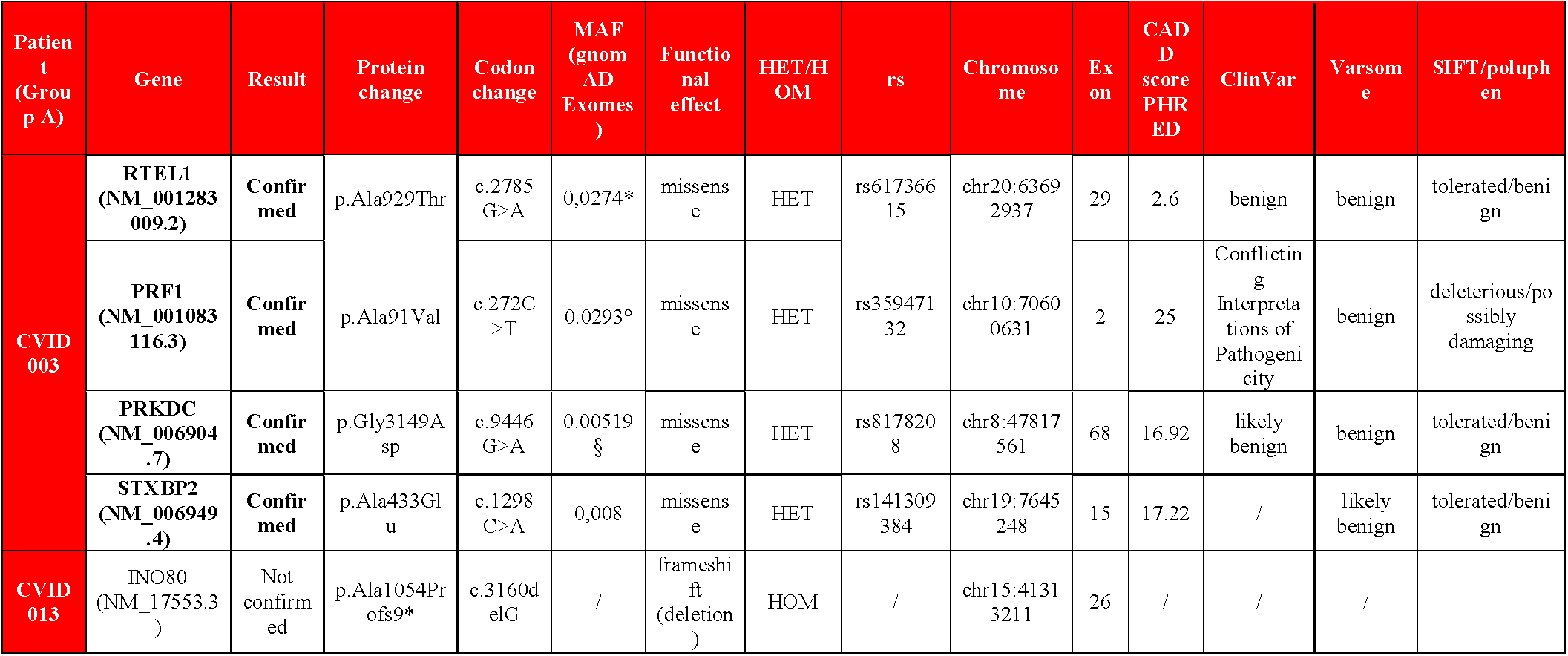
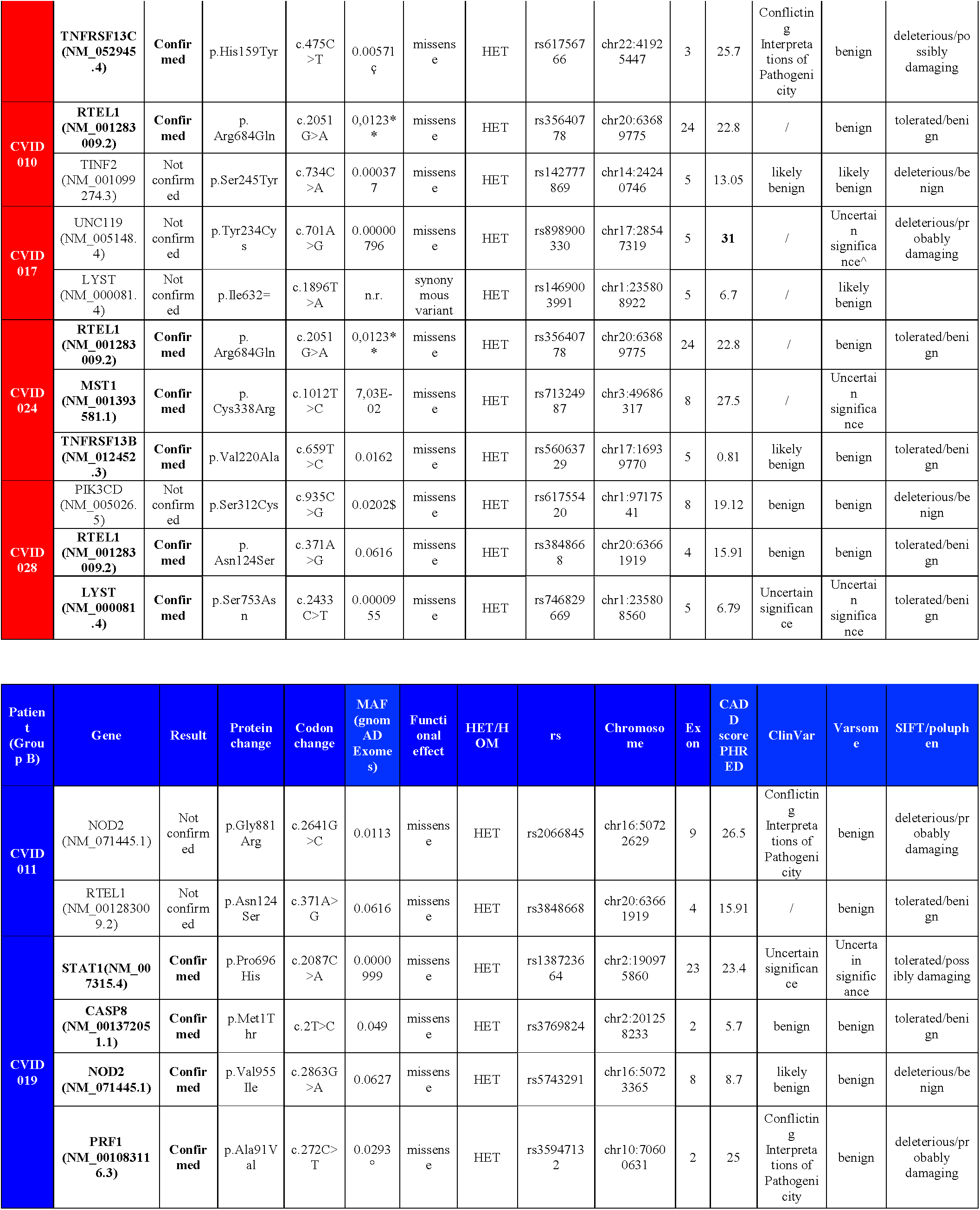
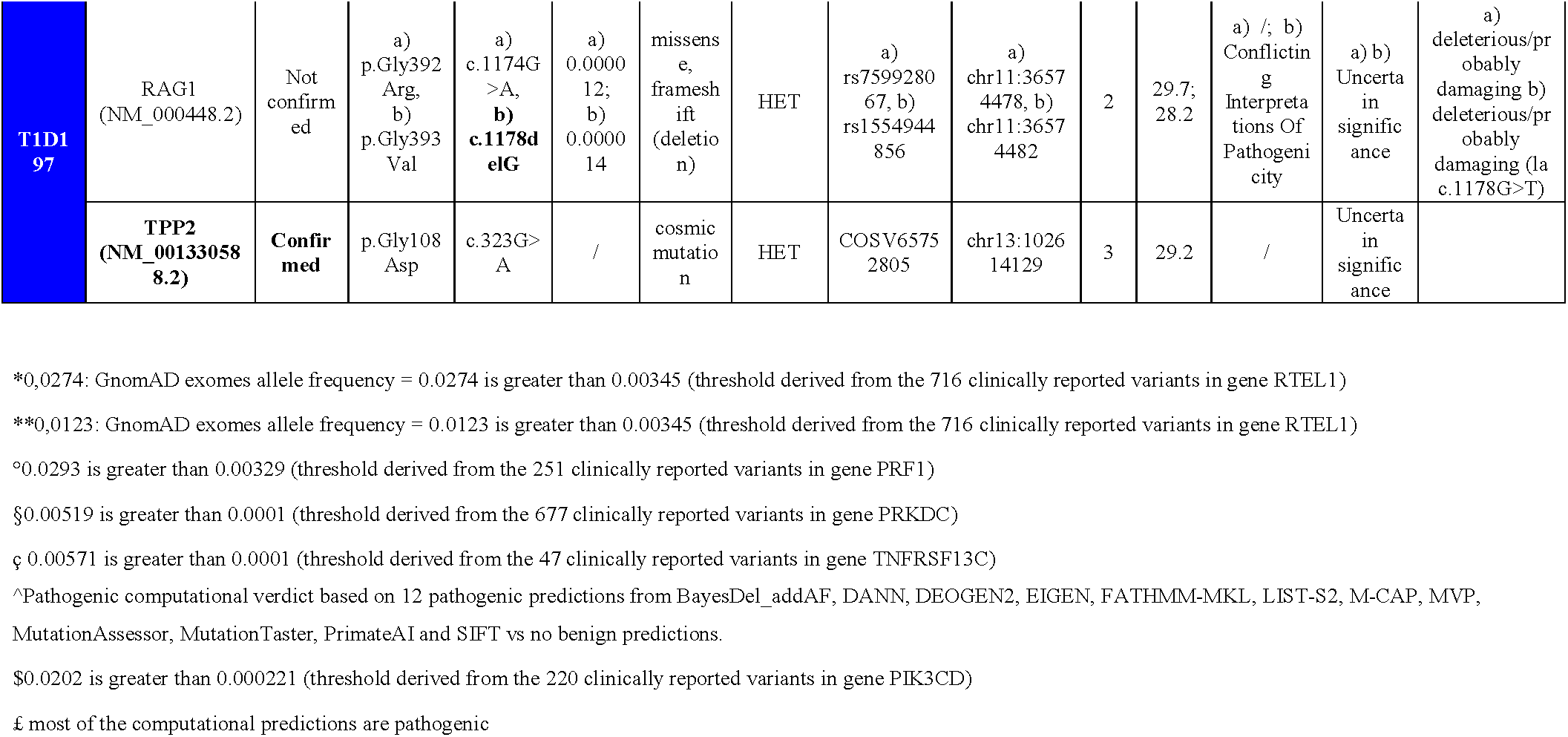
Mutations sequenced with Sanger most likely associated with CVID

RTEL1 deficiency has recently been described as the major autosomic recessive etiology of dyskeratosis congenita (DKC), a rare disease that results from excessive telomere shortening and includes bone marrow failure, mucosal fragility, pulmonary or liver fibrosis, early onset inflammatory bowel diseases, neurological impairment and, in more severe cases, immune deficiency and increased susceptibility to malignancies (36–38). Accordingly, we evaluated telomere length in lymphocyte subsets isolated from two group A patients CVID003 and CVID010, bearing respectively p.Ala929Thr and p.Arg708Gln *RTEL1* heterozygous missense variants. Patient-derived lymphocytes had significantly shorter telomeres as compared to average control (Fig 5*A-B*). CD45RA^+^ naive T cells, CD45RA^-^ memory T cells, CD20^+^ B cells, and CD57^+^ NK cells exhibited shorter telomeres compared to control (Fig 5*A-B*).

**FIGURE 5.**
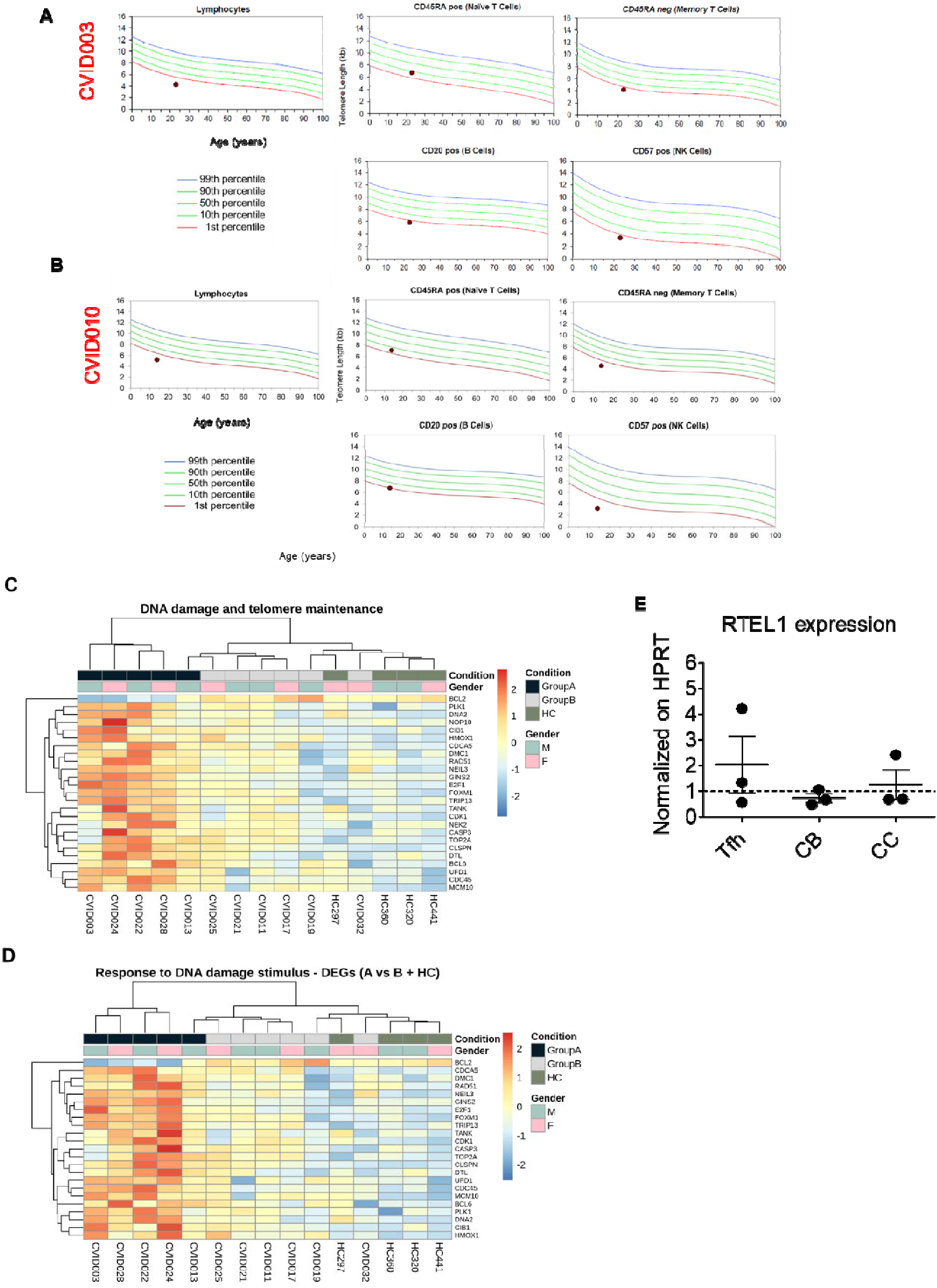
Telomere shortening in CVID patients. **(A-B)** Nomogram of Telomere Length (TL) from two CVID patients of group A, with percentile lines as annotated. TL has been measured in lymphocytes, CD45RA^+^ naive T cells, CD45RA^-^ memory T cells, CD20^+^ B cells and CD57^+^ NK. Black circle represents CVID patients. Red, green, and blue curves representing expected telomere length for the indicated proportion of healthy controls. **(C-D)** Hierarchical grouping analysis of genes involved in telomere elongation and DNA damages pathways divide patients into group A and group B. Red color intensities represent a higher gene expression. **(E)** *RTEL1* expression was assessed in sorted Tfh cells, CBs, and CCs (*n* = 3). The average for technical duplicates was estimated, normalized on *HPRT* as housekeeping gene, and represented as dark circles; *HPRT* expression (set at 1) is represented by the dotted line; mean and SD are also shown.

Next, we sought to re-anayze the RNA-Seq data and perform a biased hierarchical grouping analysis focused on DNA damage and telomere length pathways. Higher expression of genes involved in DNA damage, telomere maintenance and response to DNA damage were observed in Tfh cells from group A as compared to group B CVID patients (Fig 5*C-D*). A first comparison of RNAseq data of *RTEL1*, *TINF2*, *DKC1*, *TERT*, and *TERF1* expression levels, genes associated with DKC and Hoyeraal-Hreidarsson syndrome (HHS) (38–41), showed no expression in Tfh cells (data not included) from both CVID patients and HC. However, we further investigated *RTEL1* expression in sorted tonsillar GC Tfh cells, centroblasts (CBs) and centrocytes (CCs) from control donors (non-CVID) by ddPCR to assess if this gene was expressed within the GC.We observed that *RTEL1* expression in the GC is present and comparable to HPRT, used as housekeeping gene (Fig 5E). Taken together, genetic analyses revealed the presence of heterozygous variants in a common gene, *RTEL1*, in four CVID group A patients that clinically had in common splenomegaly and lymphadenopathy. Further analyses revealed the presence lymphocytes with short telomeres suggesting acceleration of replicative senescence.

### Splenic germinal center architecture in a patient with group A immunophenotype and heterozygous variant in *RTEL1*

Patient CVID003 with a heterozygous variant in *RTEL1*, having short lymphocyte telomeres and a group A Tfh cell immunophenotype developed splenomegaly (30 cm in maximum diameter) and underwent splenectomy. Macroscopic examination evidenced well-retained red pulp and pinpoint white pulp. Splenic parenchyma showed mild and plurifocal expansion of the white pulp with mild and focal congestion of the sinuses of the red pulp. In the subcapsular area, focal giant cell reaction of the foreign body type was associated with hemosiderin deposits (consistent with so-called ‘Gandy-Gamma’ nodules) (Fig *E6*). In the white pulp, only focal reactive GCs (Bcl-2 negative and high Ki-67 in centrofollicular cells) were seen containing an increased number of Tfh cells (CXCL13^+^ PD-1^+^), sometimes alternating with others with atrophic appearance (Fig 6*A* and data not included). The marginal zone (IgD^+^) was preserved (Fig *E6*). In the interfollicular white pulp, CD4 T lymphocytes (CD3 +, CD4 >> CD8) prevailed (data not included).

**FIGURE 6.**
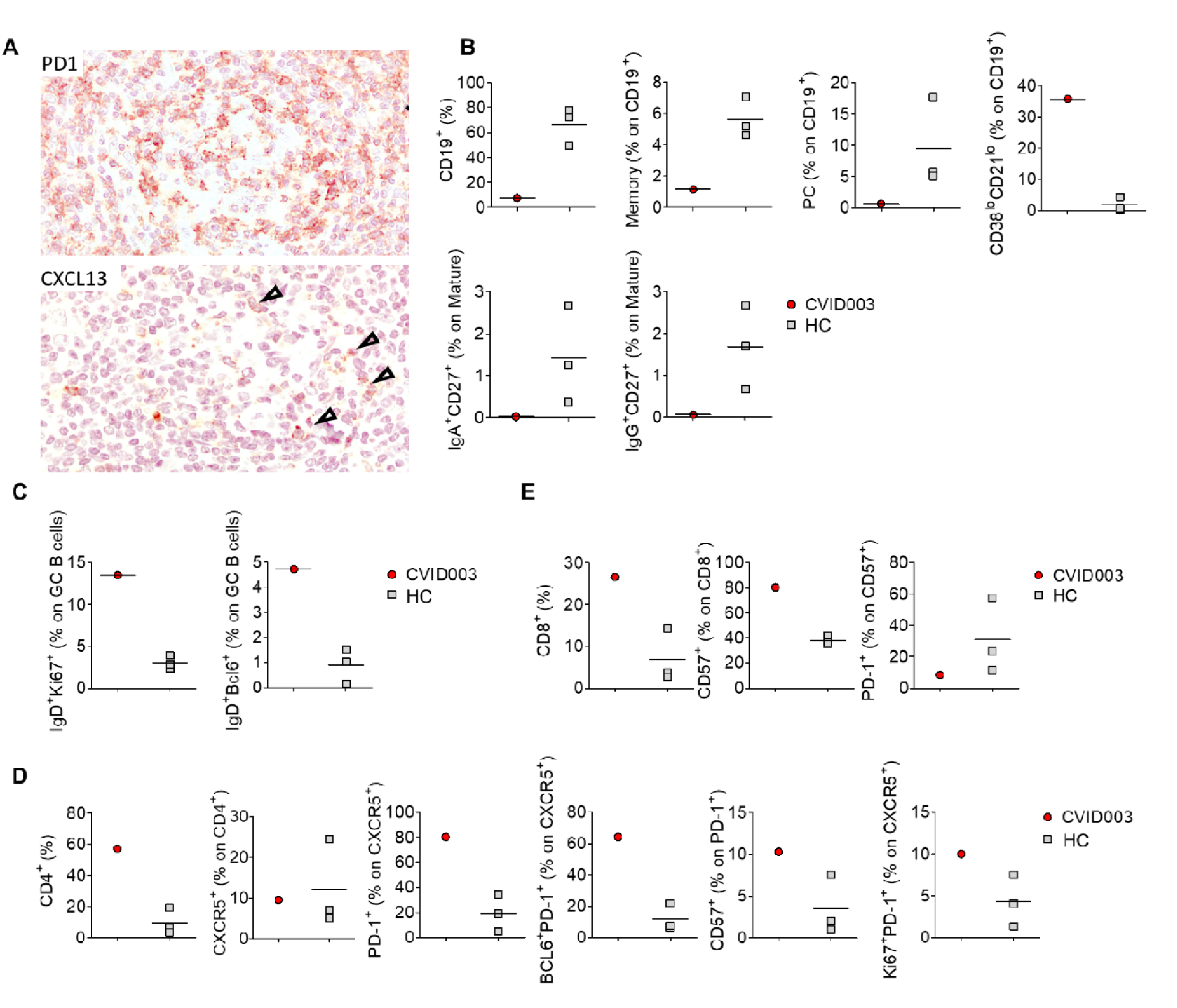
Active germinal center in the patient’s spleen. **(A)** Spleen histhopatology of CVID003 patient with RTEL1 mutation. GCs revealed an increased number of Tfh as evidenced by CXCL13 (arrowhead) and PD-1 staining. Original magnification 400X. **(B)** Percentages of CD19^+^ B cells and their subsets: memory (CD19^+^CD27^+^) B cells, CD21^lo^ B cells, plasma cells (CD38^+^CD24^-^), transitional B cells (CD38^hi^CD24^hi^), IgA^+^CD27^+^, IgG^+^CD27^-^, IgG^+^CD27^+^. **(C)** Frequencies of GC B cells expressing the proliferation markers as Ki67 and Bcl-6. **(D)** Percentages of CD4^+^CXCR5^+^ Tfh, and Tfh expressing PD-1, CD57 and Bcl-6 and Ki67 as proliferation marker in the spleen of CVID003 patient compared to age-matched controls. Percentages were analyzed by flow cytometry.

In line with the histological data, flow cytometry analysis of the splenic B cell subpopulation revealed a decrease in CD19^+^ total B cells (7.5% in CVID003 vs median 66.43% in HC), and a reduction in memory B cells (1.15% in CVID003 vs median 5,62% in HC) and plasma cells (0.62% in CVID003 vs median 9.44% in HC) as compared to control (Fig 6*B* and *E7*, for additional B cell subset analysis). Percentage of autoreactive, CD21^lo^CD38^lo^ and transitional B cells were increased in the patient (35.7% in CVID003 vs median 1.82% in HC) (Fig 6*B* and *E7*). No IgA^+^ or IgG^+^ B cells could be detected (Fig. 6*B*). Furthermore, an increase in Ki67, a proliferation marker (13.5% in CVID003 vs median 3.04% in HC) and Bcl-6 (4.72% in CVID003 vs median 0.9% in HC) expression in B cells was observed (Fig 6*C*).

Splenic CD4 and CXCR5^+^ Tfh cell frequencies were also higher in the patient as compared to control spleen (57.2% in CVID003 vs median 9.78% in HC and 9.57% in CVID003 vs median 12.1% in HC, respectively) (Fig 6*D* and *E8*), and expressed PD-1, Bcl-6, and CD57 at higher levels than the controls (PD-1: 80.20% in CVID003 vs median 19.32% in HC; Bcl-6: 64.10% in CVID003 vs median 11.96% in HC;CD57: 10.30% in CVID003 vs median 3.51% in HC) (Fig 6*D* and *E8*). Ki-67 was also highly expressed by CVID003 Tfh cells (10% in CVID003 vs median 4.31% in HC) (Fig 6*D* and *E8*). We also observed higher frequency of CD8 T cells (26.60% in CVID003 vs median 6.91% in HC) that also enriched in CD57^+^ cells (80.30% in CVID003 vs median 37.73% in HC). However, PD-1 expression levels were not increased in CD8 T cells in the spleen of the patient (8.34% in CVID003 vs median 30.87% in HC) (Fig 6*E* and *E9*).

## Discussion

This study unravels a group of CVID patients that markedly differ from usual CVID patients and HC in their Tfh cell composition. Patients within this group showed a Tfh1^hi^Tfh17^lo^PD-1^hi^CXCL13^hi^ immunophenotype, a high Th1 plasma cytokine and chemokine polarization, and a Tfh-cell RNA signature consistent with highly activated but exhausted and apoptotic cells. Equally important, genetic analysis identified monoallelic variants and polymorphisms in *RTEL1,* a helicase essential in DNA metabolism, in four patients belonging to this group, whose lymphocytes presented significantly shortened telomeres. These results were achieved by evaluating a broad array of GC-related immune markers in the blood of a subset of CVID patients presenting AI as a secondary complication exploiting a multifaceted investigative approach including flow cytometry, genome sequencing and transcriptomic evaluation of Tfh cells. Hence, our findings indicate that heterozygous variants in DNA damage response and telomere elongation pathways could underlie CVID and be strongly linked to GC-associated immune dysregulation.

CVID is a collection of disorders of humoral dysregulation resulting in low IgG and Ig/IgM levels and antibody-specific responses with recurrent infections (43). Several previous studies have addressed the immunologic components of CVID in the peripheral blood and tissues of the affected patients and it showed that the B cell and Tfh cell profile are able to determine different forms of the disease (11). Our flow cytometry and RNA seq analyses of Tfh cells separated our cohort of CVID patients into two distinct groups, one of which was characterized by a pronounced splenomegaly and lymphadenopathy. Our data suggest that this approach can be used to identify patients with a higher risk for immune-related dysregulation, shortened telomeres, and perhaps those carrying defects in DNA synthesis and repair. We found that the Tfh cell signature (high levels of PD-1 and Th1 polarization) correlated with plasma CXCL13 levels and CD21^lo^CD38^lo^ autoreactive B cells, suggesting the use of CXCL13 as an additional potential biomarker for this form of CVID. Additional studies in larger cohorts of patients will be required to establish the set of biomarkers that define this form of CVID and direct future therapeutic lines of research.

Tfh cells are pivotal players during the GC reaction and the production of high affinity, long-lived antibody responses (44). In our CVID patients Tfh cells retain B cell helper activity as evidenced by their ability to promote plasmablast differentiation and IgM production, suggesting that the defect in Ig class-switching was rather B-cell intrinsic. This notion is corroborated by our RNA seq data showing an overall normal Tfh cell signature in group B patients and a rather hyperactive Tfh signature in those belonging to group A. We cannot exclude, however, that the Th1 phenotype of Tfh cells played a role in preventing efficient GC responses and class switching, as the addition of IFNγ was shown to reduce IgG and IgA production in T/B co-cultures (11). There is also evidence from previous clinical settings, i.e., HIV and CVID, that Tfh1 cells are less effective B cell helpers in comparison to their CXCR3^-^ Tfh counterparts (45–47). Interestingly, a predominant Tfh1 cell immunophenotype has been detected in several autoimmune diseases and syndromes (33,48), suggesting that excessive IFNγ production in GC might promote AAb production.

RTEL1 has been proposed to dismantle T-loops during replication thus preventing catastrophic cleavage of telomeres as a whole extra-chromosomal T-circle (49). Previous observations indicated that heterozygous *RTEL1* mutations are associated with premature telomere shortening despite the presence of a functional wild-type allele *in vivo* (36,50). Furthermore, Speckmann *et al* showed that the immunological and clinical phenotype is very much mutation/variant-dependent but, overall, premature telomere shortening is a common feature (36). Our group of patients with RTEL1 variants exhibited very short telomeres in their lymphocytes. Tfh cells expressed genes involved in DNA synthesis and repair, telomere maintanance, apoptosis, and exhaustion. Furthermore, we detected RTEL1 expression in tonsillar germinal center Tfh cells. We therefore assume that the observed CVID phenotype in group A patients with RTEL1 variants a may be a consequence of Tfh-cell replicative senescence and exhaustion upon repeated proliferation stimuli triggered by pathogens.

RTEL1 expression was also detected in GC B cells. Tonsil-isolated centrocytes and centroblasts expressed similar levels of RTEL1. Interestinlgy, V(D)J recombination efficiency in RTEL1 deficiency(36) was previously found unaffected and comparable to HC suggesting a normal B-lymphocyte development in the germinal centers and, possibly regular production of PCs. Possibly, RTEL1 mediates proliferative senescence also in mature B cells due to its essential role in DNA replication, homologous recombination, and telomere maintenance. (51). Due to its well-documented role in CD34^+^ hematopoietic stem cells (HSC), RTEL1 deficiency my have contributed to B cell failure and hypogamaglobulinemia via proliferative exhaustion of the HSC compartment. In future studies we hope to address whether the bone marrow compartment was affected in our patietns with variants in *RTEL1*.

A previous study reported short telomers and reduced capacity to divide in T and B cells from a subset of patients with CVID (52). It would be interesting to address whether Tfh cells and CXCL13 levels are similar to our group A patients and if variants in RTEL1 or other associated genes can be found. Of interest, Bcl-6, the master regulator of the GC B and Tfh cell lineage, is located on chromosome 3q27 at the telomere proximity (53). RTEL1 is also located at the telomere proximity of chromosome 20. Recent studies have suggested that telomere length regulates the expression of genes that are located up to 10 Mb away from the telomere long before telomeres become short enough to produce a DNA damage response (senescence) (54)(55). This suggests that excessive telomere shortening could have played a role on immune cell fitness through another and till today unexplored mechanism leading to this form of CVID.

Analysis of the spleen in one patient revealed high frequency of CD57^+^ and PD-1^+^ CD4 and CD8 T cells. CD57 is a marker of GC Tfh cells (56) but also a marker of T-cell replicative senescence associated with short telomeres (57). CD57^+^ T cells are characterized by an inability to undergo new cell□division cycles despite preserved ability to secrete cytokines after antigen encounter (57,58). Interestingly, Klocperk et al identified a population of follicular CD8 T cells in the lymph nodes of patients with CVID who clinically were characterized by lymphadenopathy. These follicular CD8 T cells also displayed senescence-associated (CD57) features suggesting they were exhausted(59). Group A patients showed high levels of cytokines involved in inflammatory response in their serum (e.g., INF-γ, IL-2R, MDC, MIP3-α, SDF-1α), suggesting that their CD4 (Th and Tfh) and perhaps CD8 T cells were able to produce cytokines despite their exhaustion phenotype.

In conclusion, by characterizing the phenotype and transcriptome of circulating Tfh cells in patients with CVID, we were able to identify a group of patients with specific clinical and immunological characteristics most possibly influenced by the presence of pathogenic variants/polymorphisms in *RTEL1*. Despite the limitation of a very small sample size, our data suggest that a Tfh1^hi^Tfh17^lo^PD-1^hi^CXCL13^hi^ immunophenotype and short lymphocyte telomere length could be used as indicators for genetic testing of *RTEL1* and possibly other DKC causing genes (*TERT*, *DKC1*, *NHP2*, *TERC* etc) in patients with CVID. Further studies will be required to better understand the contribution of *RTEL1* in Tfh and B cell development, function and interactions, and whether the alterations seen in CVID patients with *RTEL1* variants are genetically-driven and/or secondary to infections and chronic immune stimulation.

## Abbreviations

CVID: Common variable immunodeficiency
PAD: Primary antibody deficiency
HC: Healthy control
Tfh: T follicular helper
Treg: T regulatory
Tfr: T follicular regulatory
GCs: Germinal centers
CXCR5: Chemokine (C-X-C motif) receptor type 5
PD-1: Programmed cell death protein 1
ICOS: Inducible T cell co-stimulator
CXCL13: Chemokine (C-X-C motif) ligand 13
WES: Whole Exome Sequencing
Bcl-6: B cell lymphoma 6
RTEL1: Regulator of telomere elongation helicase 1
AICD: Activation-induced cell death
B_M_: B memory
B_N_: B naїve
DKC: Dyskeratosis congenita
CBs: Centroblasts
CCs: Centrocytes
HSC: Hematopoietic stem cells
SCID: Severe combined immunodeficiency
ESID: European Society for Immunodeficiencies
GSEA: Gene Set Enrichment Analysis

## Acknowledgements

This work was supported from 5×1000 OSR PILOT & SEED GRANT to GF & MPC. We would like to thank our past lab members for their contribution. Moreover, we acknowledge the nurses, the patients and their families.

## Supplementary

### Materials and Methods

#### B cell helper assay

PBMCs were sorted into CD19^+^CD38^-^CD27^-^ naive B cells, CD19^+^CD38^-^CD27^+^ memory B cells, and CD25^-^CXCR5^+^ Tfh cells. Prior to sorting, PBMCs were stained with a panel of mAbs that consisted of: CD19-FITC (4G7, BD Biosciences), CD27-PE (L128, BD Biosciences), CD25-APC (2A3, BD Biosciences), CD4-PE-Vio770 (M-T321, Miltenyi Biotec), CXCR5- Brilliant Violet 421 (J252D4, BioLegend), and sorted using a FACSAria Fusion sorter cytometer (Becton Dickinson). B cells (3 x 10^4^) were co-cultured with an equal number of CD25-CXCR5+ sorted Tfh cells and stimulated with Staphylococcal enterotoxin B (100 ng/mL, Sigma-Aldrich) in complete RPMI. On culture day 7, the frequency of plasmablasts CD38^+^CD20^low^ was analyzed by flow cytometry. Culture supernatant IgM and IgG concentrations were determined by ELISA assay (Human IgM and IgG Uncoated ELISA Kit, Invitrogen by Thermo Fisher Scientific, Cat. No. 88-50620-22 and 88-50550-22) according to the manufacturer’s instructions.

#### Multiplexing protein level measurements on a single Luminex platform

Secreted protein levels in sera were detected using the Invitrogen^TM^ ProcartaPlex^TM^ Human 65-plex panel kit (Thermo Fisher Scientific Cat. No. EPX650-10065-901). Samples were assayed according to the manufacturer’s instructions (60), and the plates were read on a Luminex xMAP instrument (BioRad). The acquisition and analysis of the samples were performed with the Bio-Plex Manager 6.0 software (BioRad).

## Summary

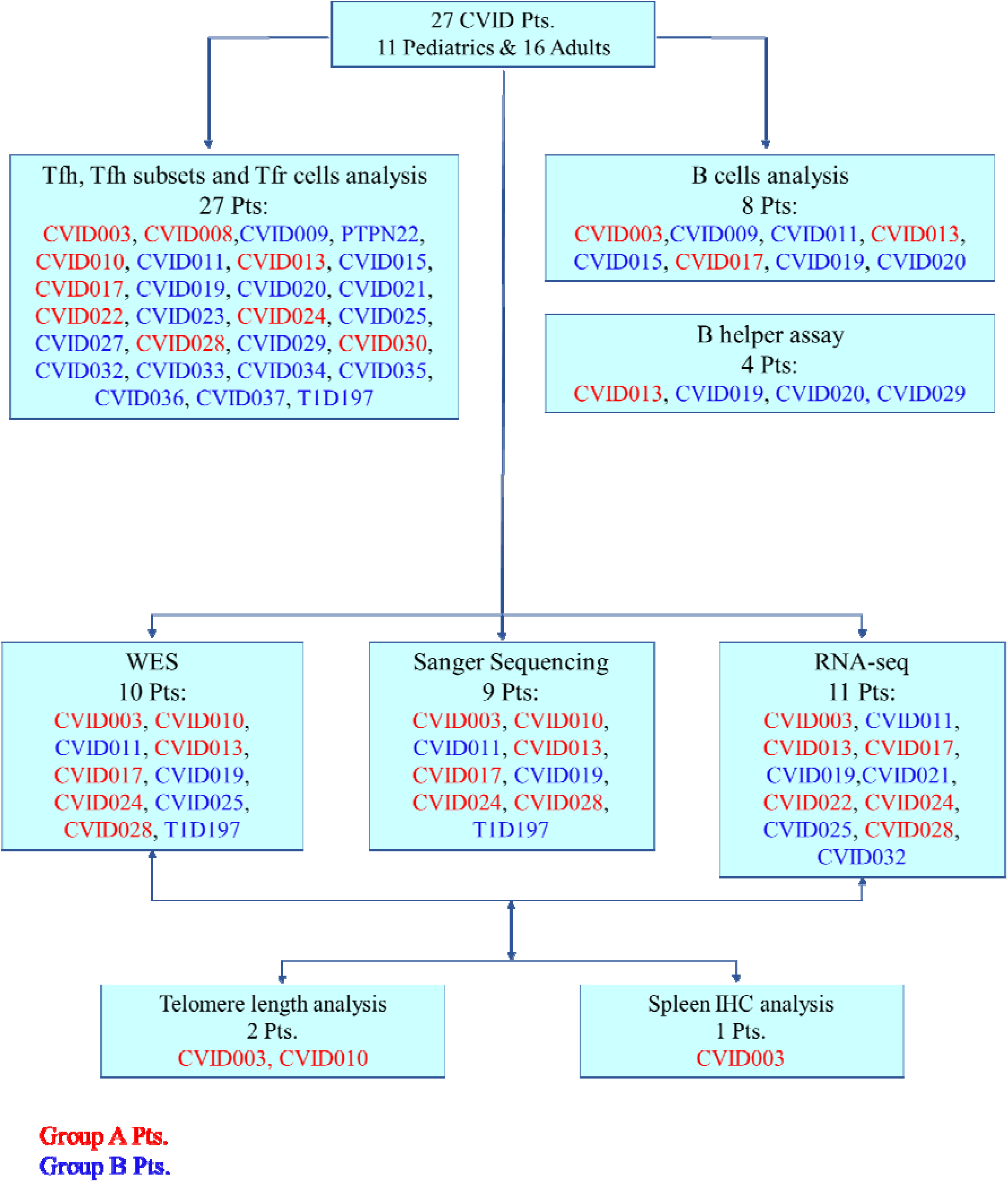

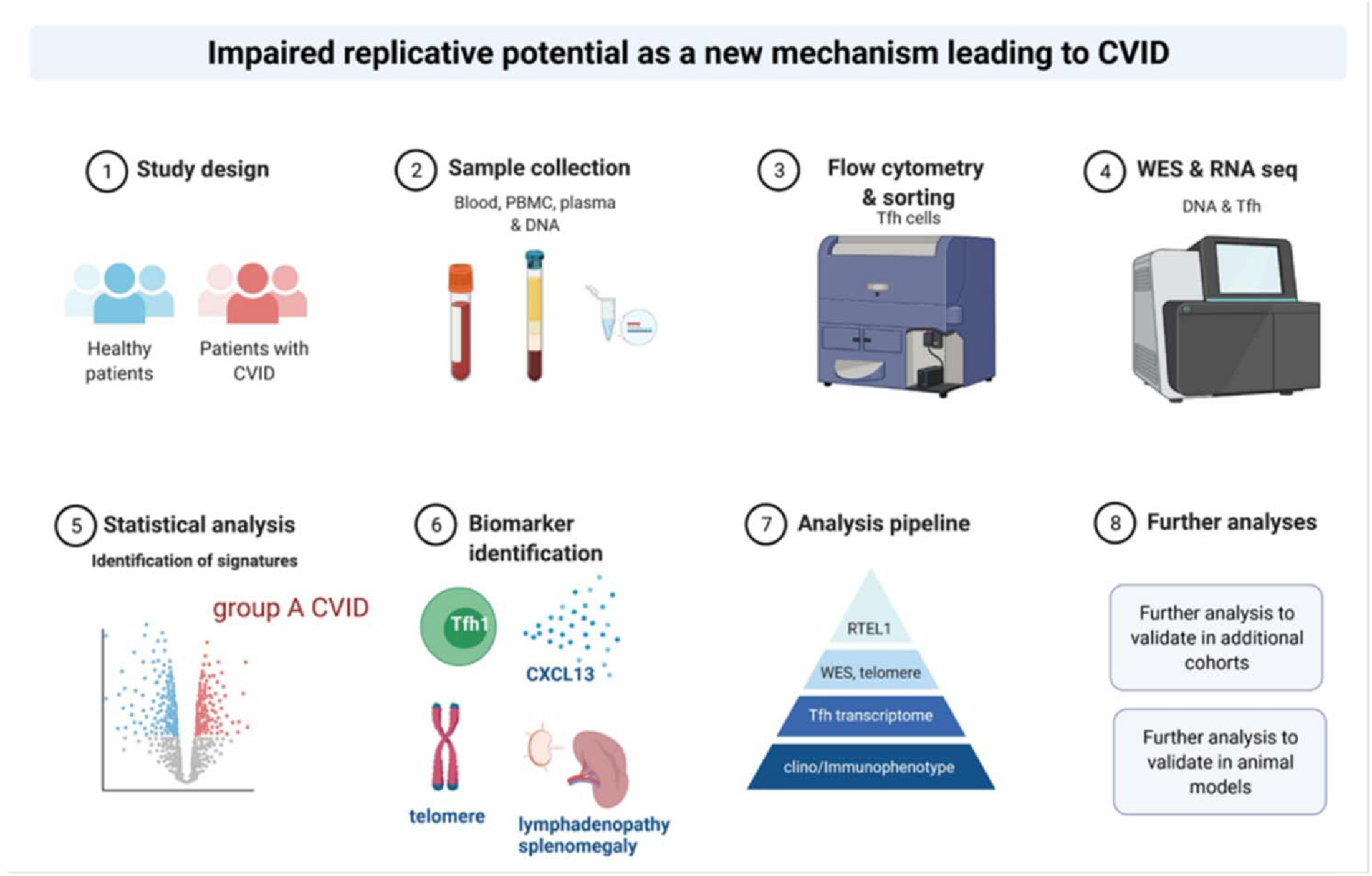

### Supplementary Figures

**FIGURE E1.**
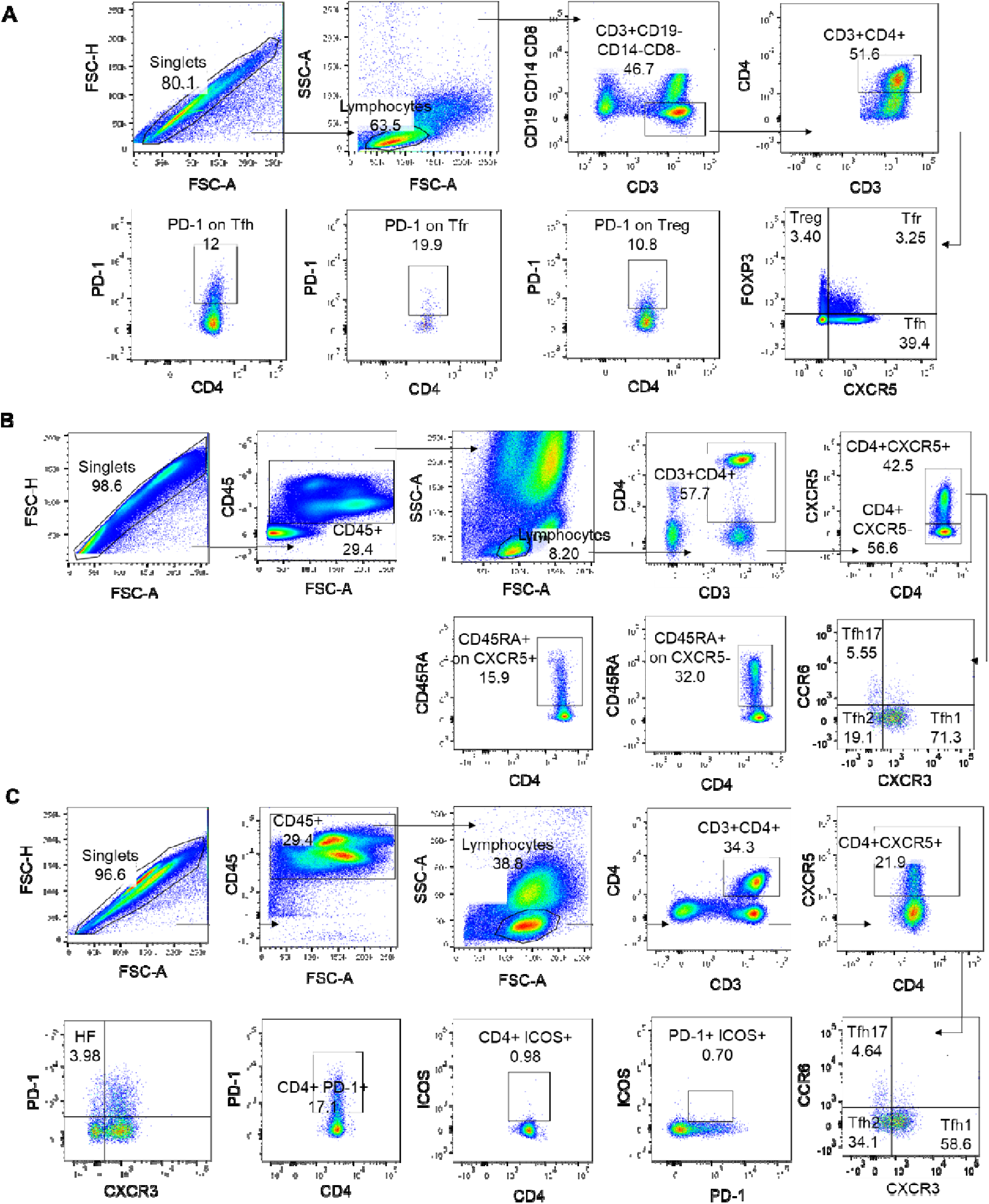
Representative gating strategy for **(A)** Tfh (CXCR5^+^FoxP3^-^), Tfr (CXCR5^+^FOXP3^+^, Treg (CXCR5^-^FOXP3^+^) cells, the activation marker (PD-1), **(B)** Tfh substets and **(C)** Highly Functional Tfh (PD-1^+^CXCR3^+^) cells.

**FIGURE E2.**
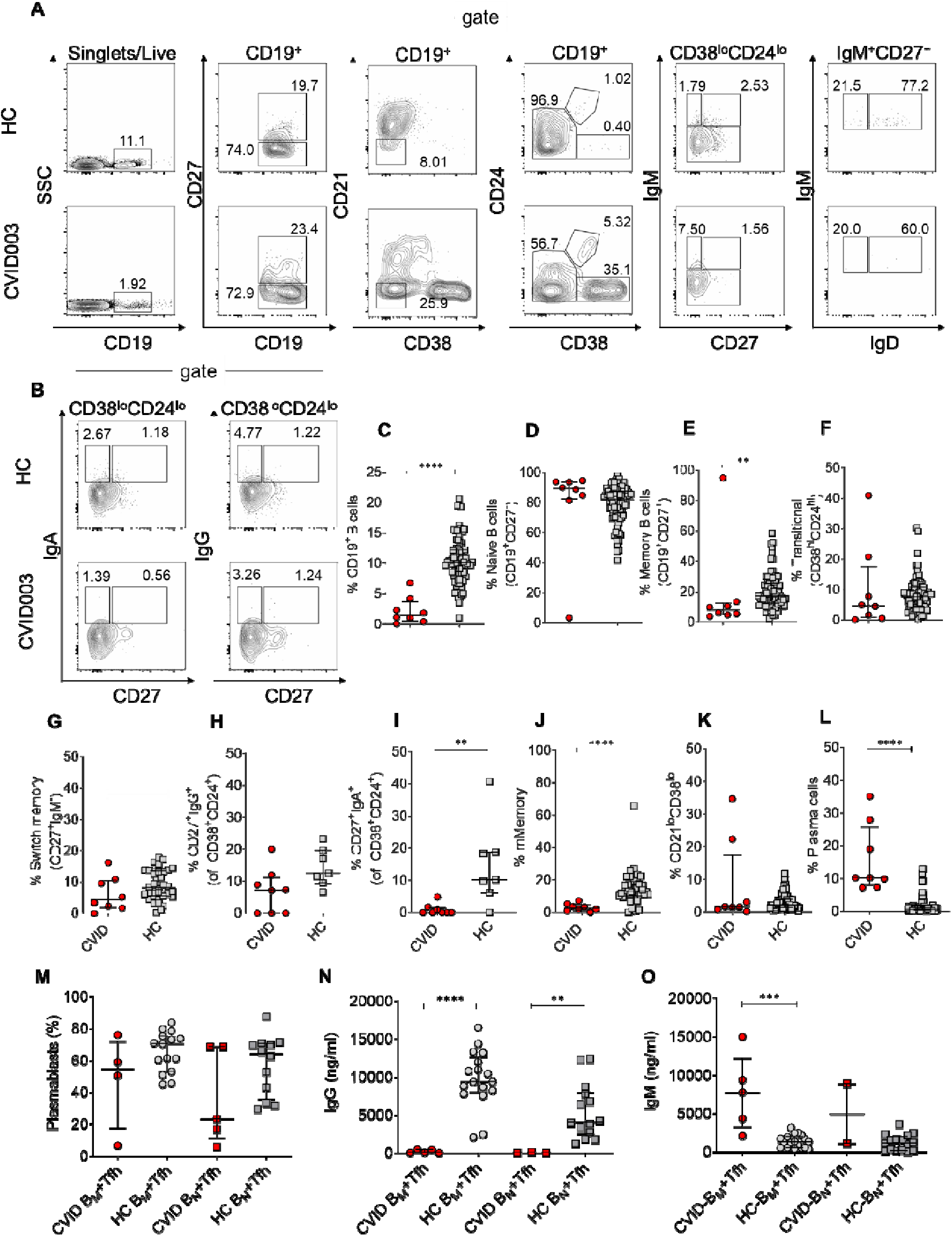
**(A)** Phenotypic characterization of peripheral blood B cells and B-cell subsets. Gating strategy to identify naïve, memory and switch memory B cells based on expression of CD24, CD38, CD27, and **(B)** IgH isotypes. **(C)** Flow cytometric analysis of B cells and B-cell subpopulations isolated from spleen of CVID003 patient with *RTEL1* mutation compared to age-matched controls. Frequency of CD19^+^ B cells, **(D)** naïve (CD19^+^CD27^-^) and **(E)** memory (CD19^+^CD27^+^) B cells, **(F)** transitional B cells (CD38^hi^CD24^hi^), **(G)** switched memory (CD27^+^IgM^-^) B cells and **(H-I)** their 2 subclasses CD27^+^IgG^+^ and CD27^+^IgA^+^. **(J)** Frequencies of mature memory B cells (CD27^+^IgM^+^), **(K)** CD21^lo^ B cells (IgM^+^IgD^+^) and **(L)** plasma cells (CD24^-^CD38^+^). **(M)** Percentages of plasmablast (CD38^hi^ CD20^-^ CD19^+^) after 1 week co-culture of B naïve (B_N_) or B memory (B_M_) cells with Tfh cells in CVID patients compared to controls. Percentages were analyzed by flow cytometry. **(N-O)** IgG and IgM measured in ng/mL by ELISA assay in the supernatant of co-culture after 1 week. Percentages were analyzed by flow cytometry. In all graphs, red points represent individual donors and asterisks indicate statistical significance as calculated by Mann Whitney test. Black bars: median with interquantile range. *p<0,05; **p<0,005; ***p<0,001; ****p<0,0001.

**FIGURE E3.**
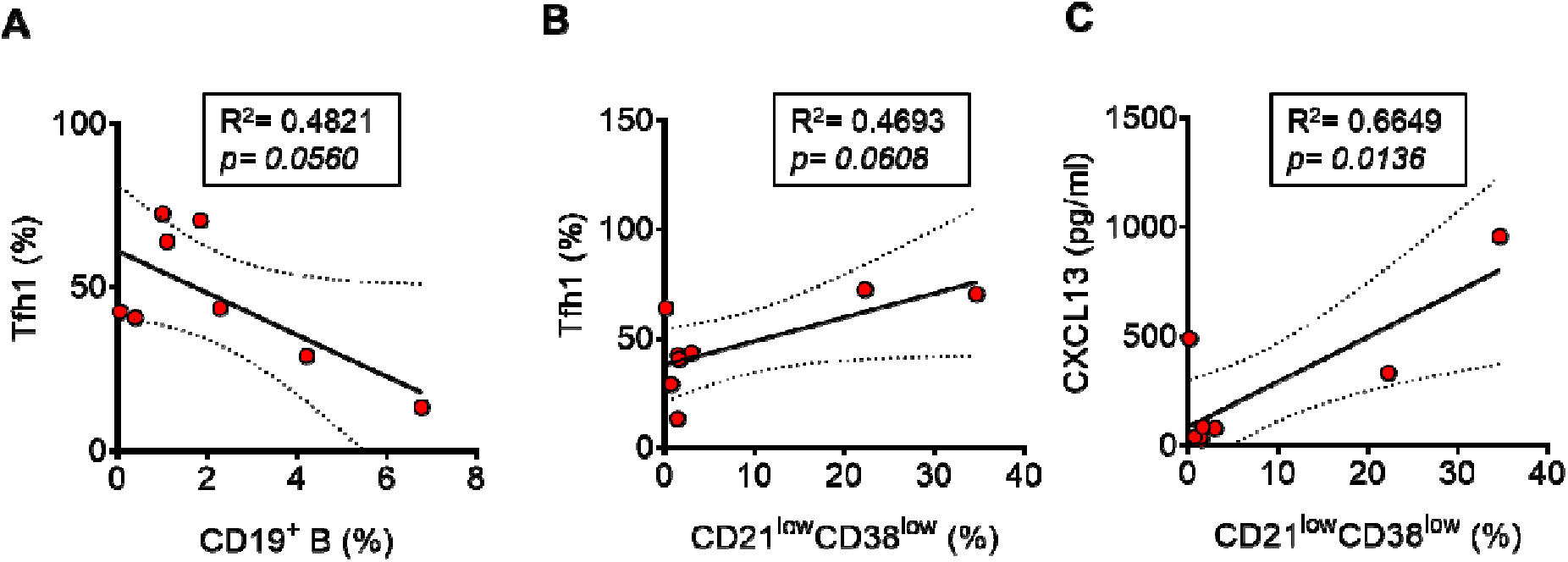
Correlation analysis between frequencies of Tfh1, CD19^+^, CD21^lo^ B cells **(A-B)** and CXCL13 plasma levels **(C)** in CVID patients. Frequencies were analyzed by flow cytometry. Lines represent linear regression and SD. *p<0,05; **p<0,005; ***p<0,001; ****p<0,0001.

**FIGURE E4.**
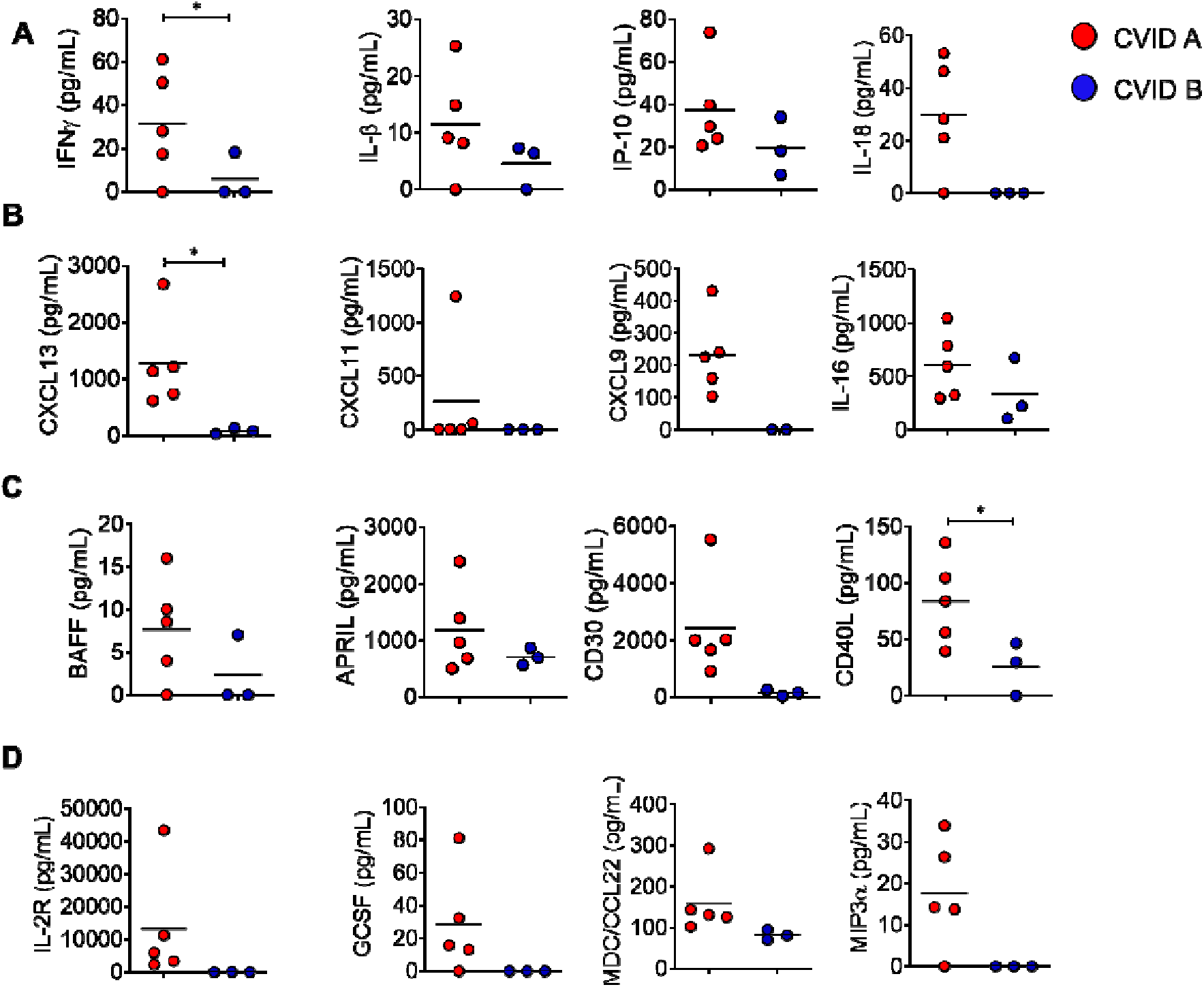
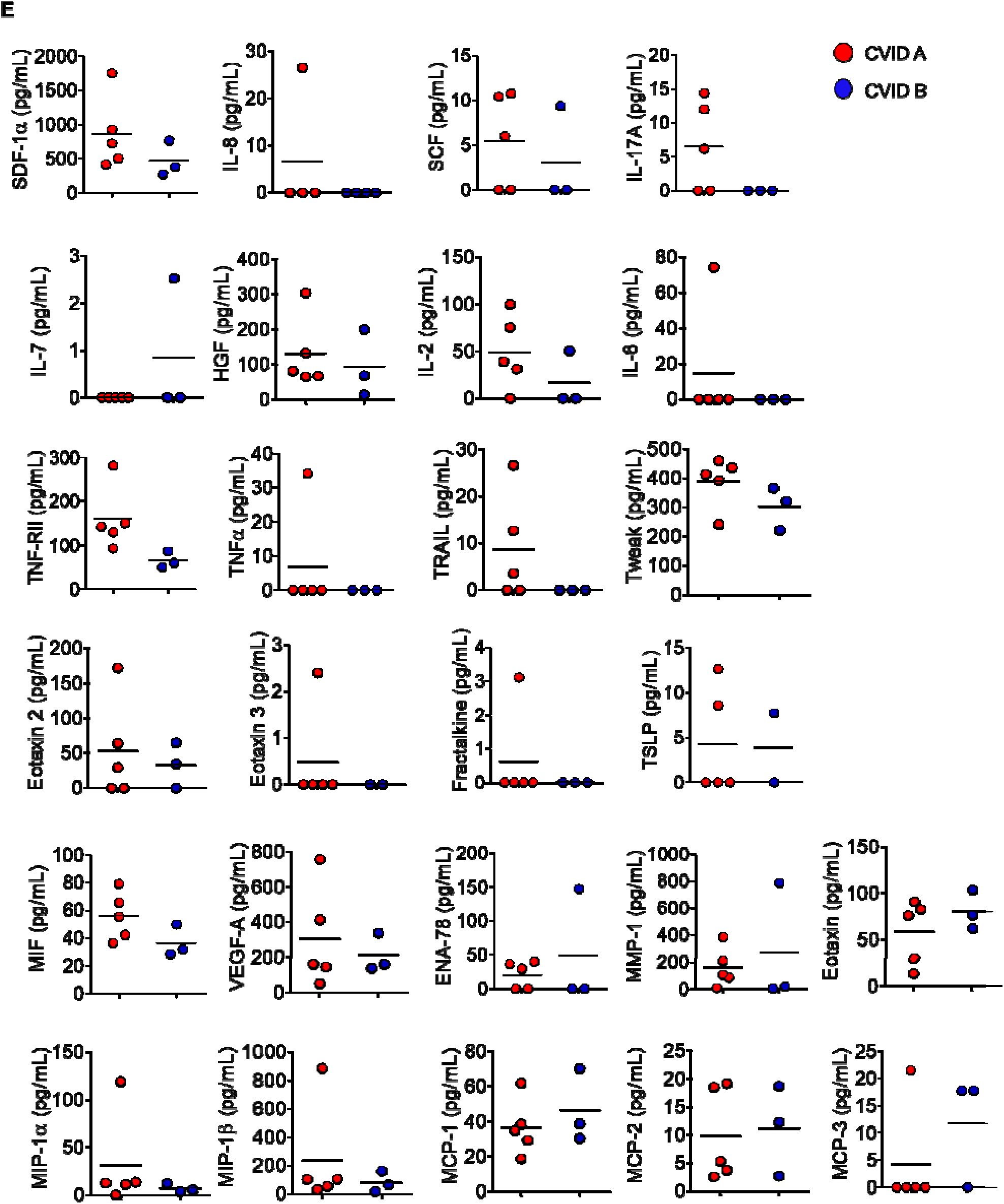
42 gene targets showed expression changes in CVID group A patients vs CVID group B. The ProcartaPlex 65-plex data was analyzed using the Bio-Plex Manager Software 6.1.

**FIGURE E5.**
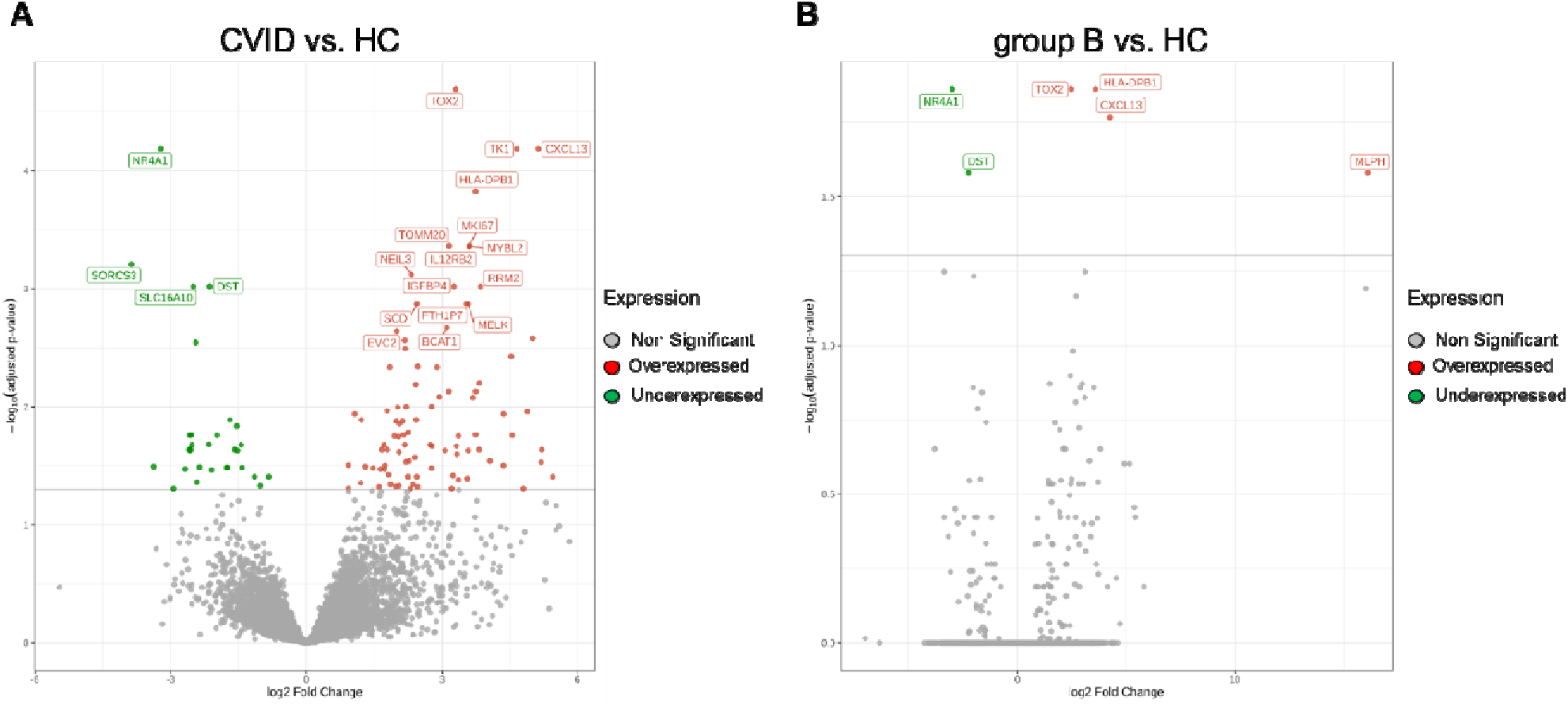
Volcano plot showing the transcriptomic analysis of sorted CD4^+^CXCR5^+^CD25^-^ Tfh cells in total CVID patient vs. HC **(A)** and in group B patients vs. healthy control **(B)**. In the volcano plots, red color represent a higher gene expression compare to green color which is for genes low expressed.

**FIGURE E6.**
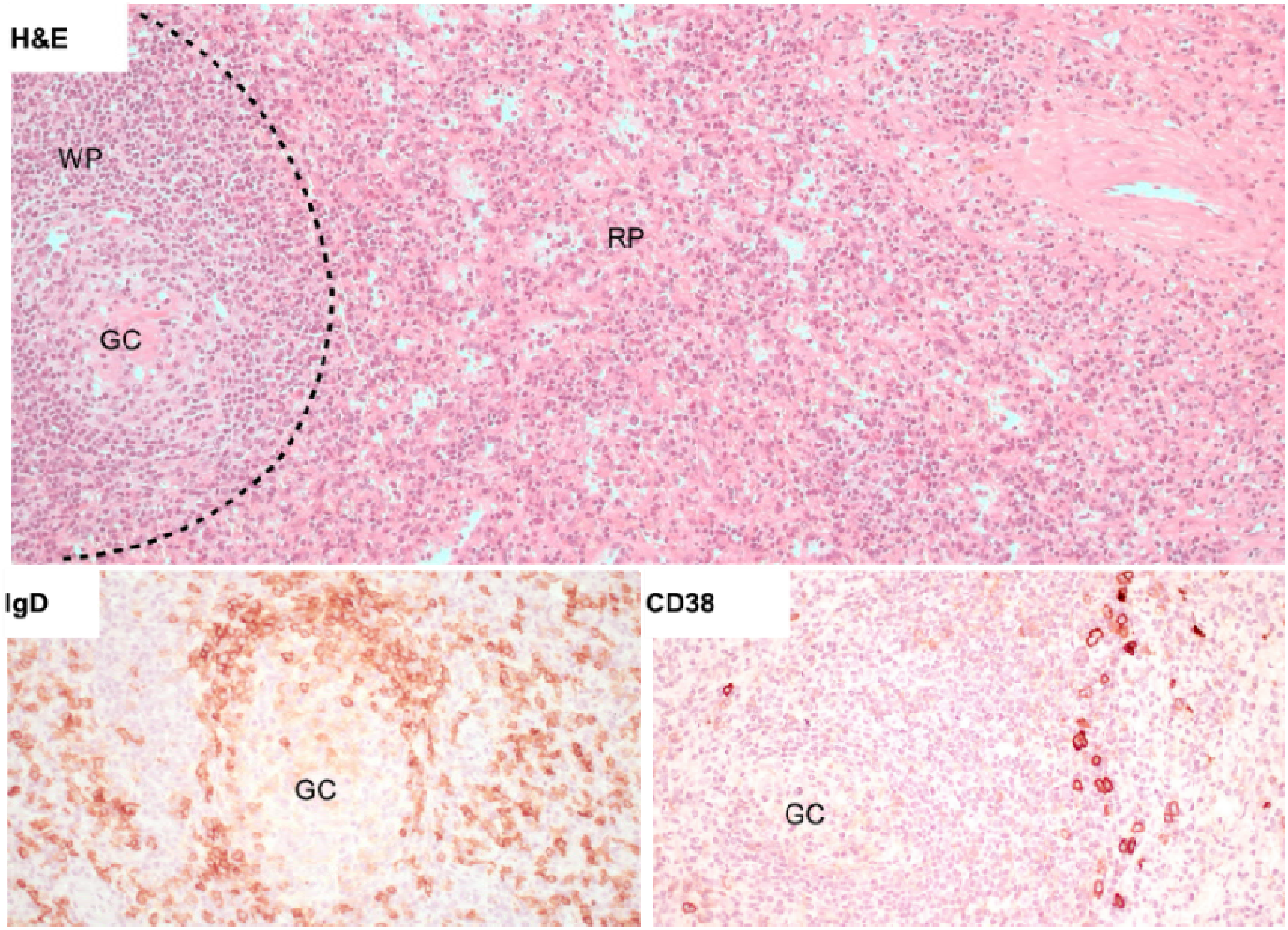
Spleen histopathology of CVID003 patient detected in formalin-fixed paraffin-embedded sections by haematoxylin and eosin (H&E) staining (above) or by immunohistochemistry staining (below) showing a reduced white pulp (WP) and down sized germinal centers (GC) while the red pulp (RP) retained a normal appearance. On immunohistochemistry a reduced marginal zone is evident (IgD); sparse plasma cells can also be detected (CD38). Original magnification: H&E 100X; IgD and CD38 200X.

**FIGURE E7.**
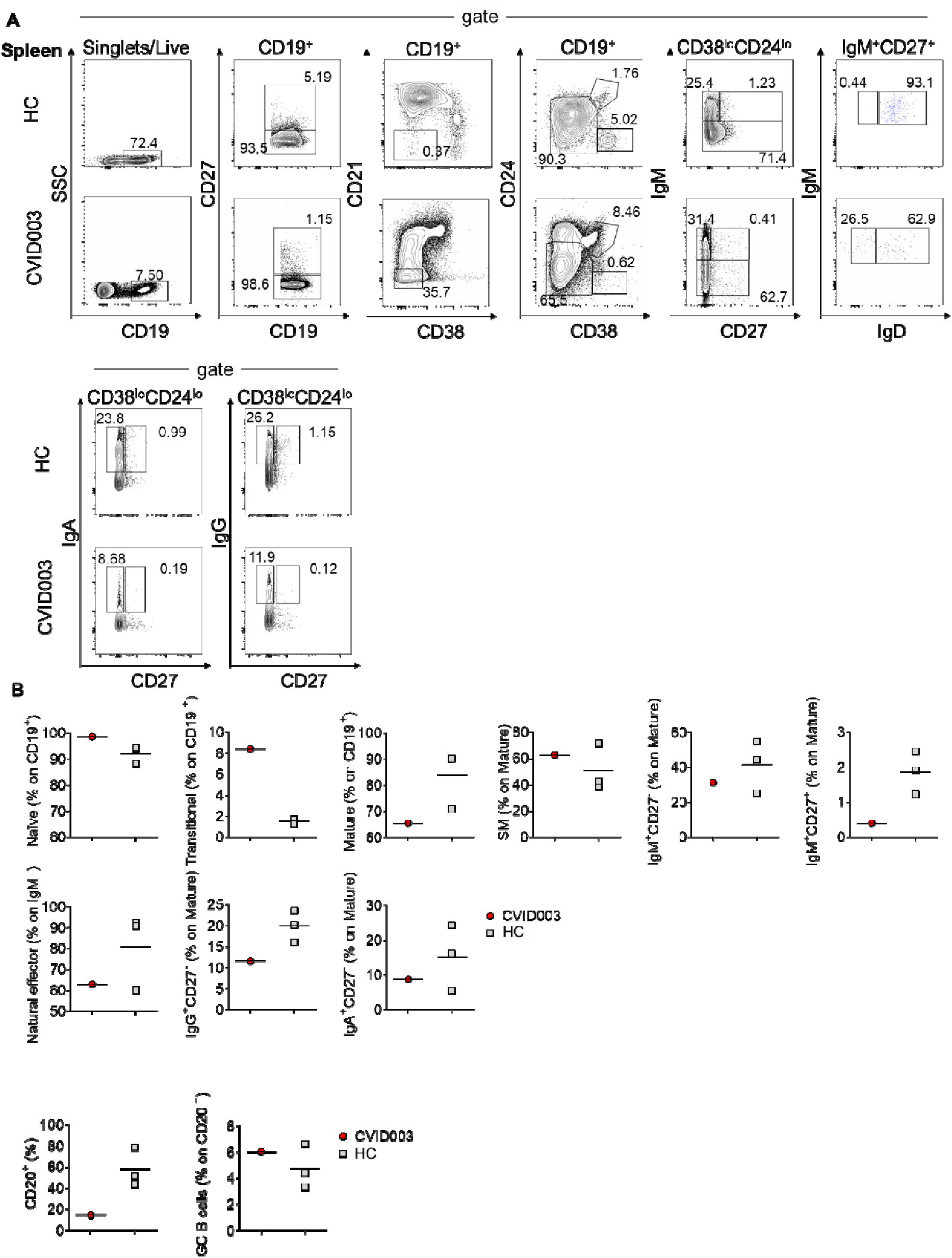
Phenotypic characterization of B cells and B-cell subsets isolated from spleen of CVID003 patient. **(A)** Gating strategy to identify naïve, memory and switch memory B cells based on expression of CD24, CD38, CD27, and **(B)** IgH isotypes.

**FIGURE E8.**
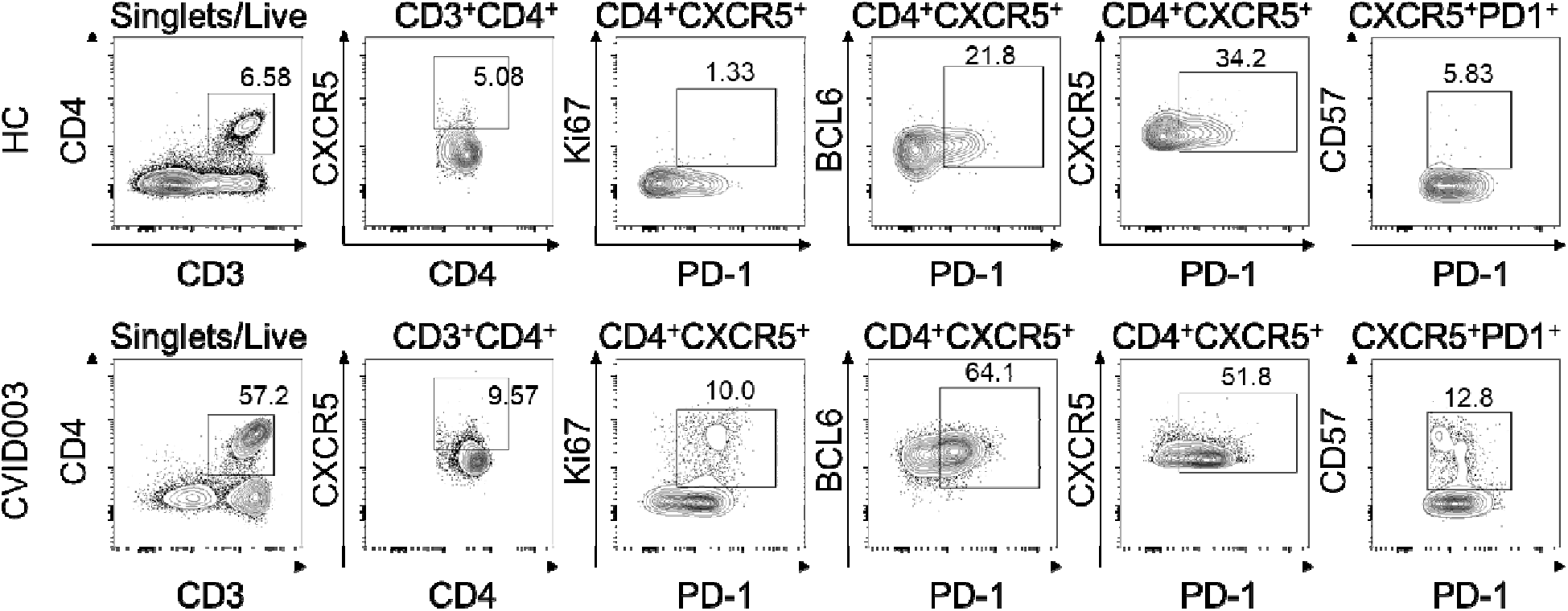
Flow cytometric analysis of CD4^+^ T cells, Tfh (CXCR5^+^CD4^+^), PD-1^+^ and CD57^+^ Tfh and germinal center’s markers (Ki67 and Bcl-6) on the spleen of CVID003 compared to age-matched controls.

**FIGURE E9.**
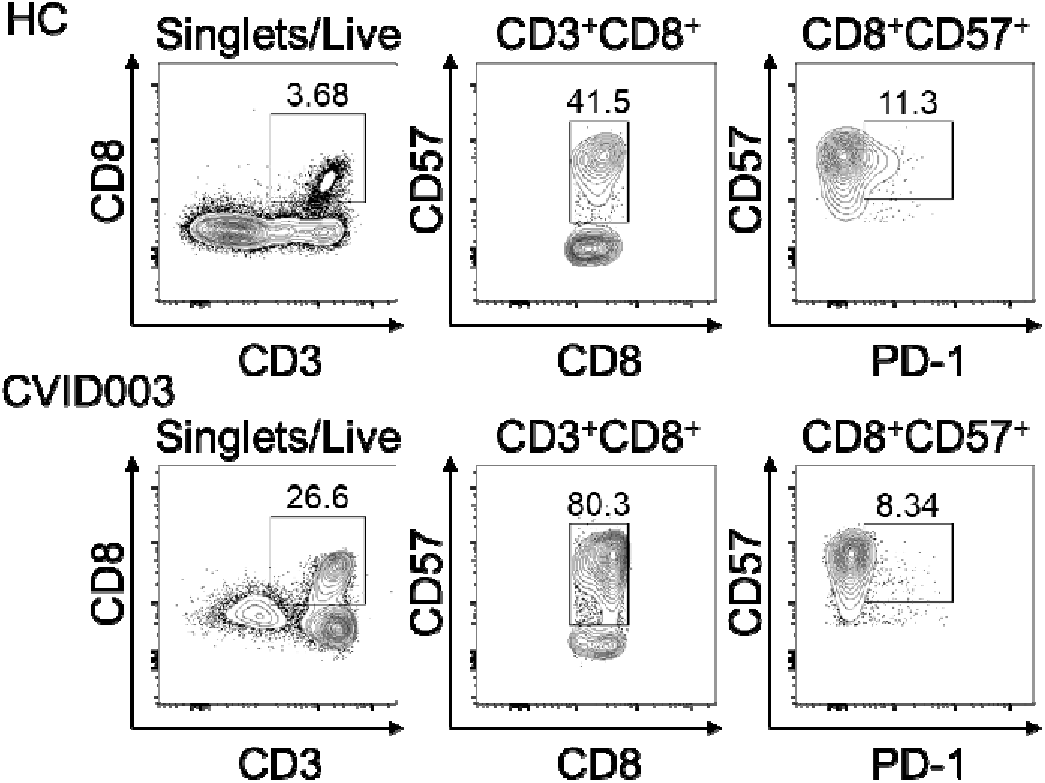
Flow cytometric analysis of activated CD8^+^ T cells isolated from the spleen of CVID003 compared to age-matched controls.

**FIGURE E10.**
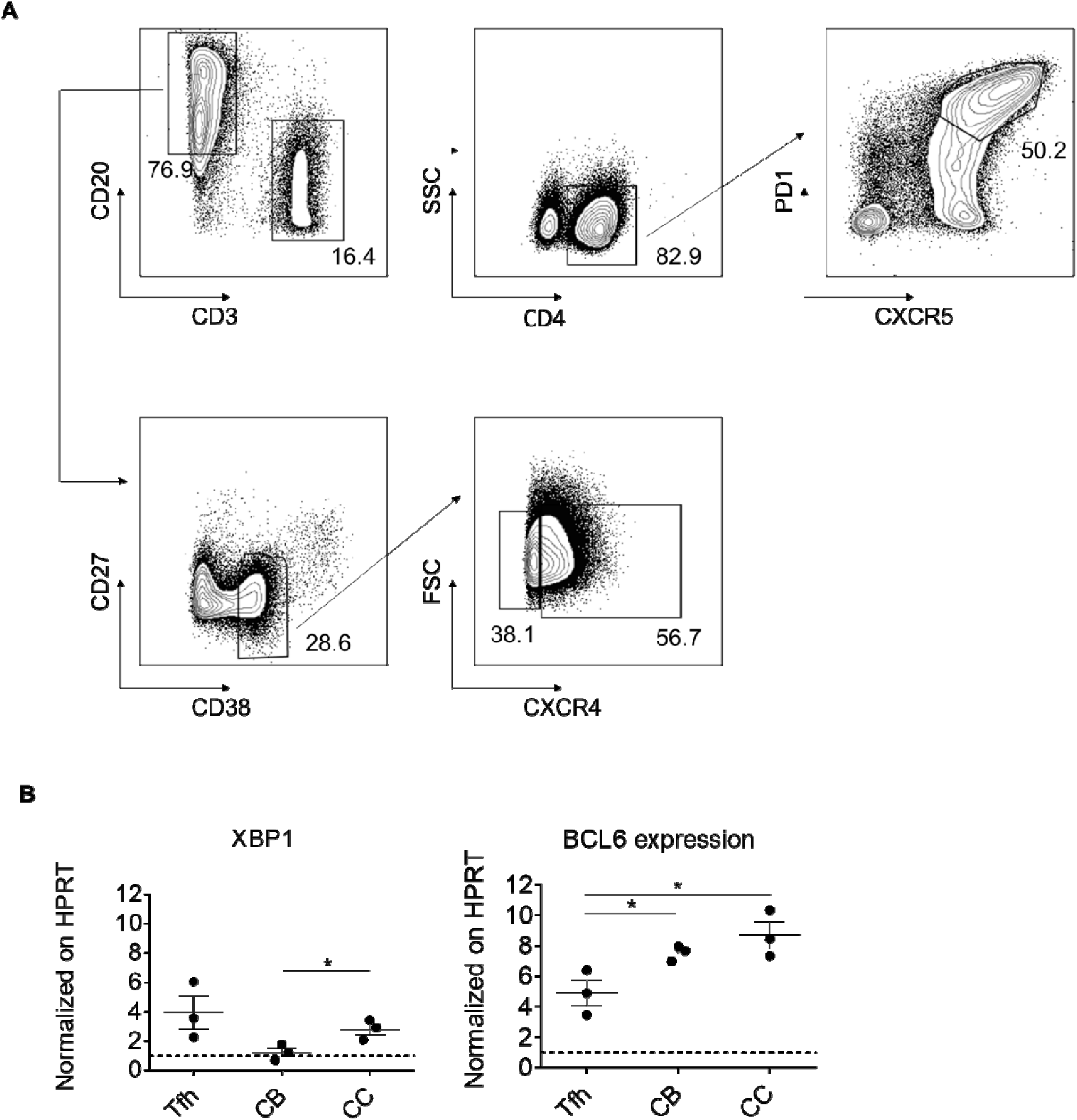
**(A).** Representative gating strategy for Tfh cells, CBs, and CCs sorting. Cells sorted as follow; Tfh cells as CD3^+^CD20^-^, CD4^+^, PD-1^+^CXCR5^+^; CCs as CD20^+^CD3^-^, CD27^-^CD3 CXCR4^-^; CCs as CD20^+^CD3^-^, CD27^-^CD38^dim^, CXCR4^+^. The complete antibody mix is described in Supplementary Table E3, panel E. **(B).** *XBP1* and *BCL6* expression was assessed in sorted Tfh cells, CBs and CCs (*n* = 3). *XBP1* was included as quality indicator for CCs and CBs sorting, since evidences from published dataset (http://biogps.org/dataset/E-GEOD-15271/) indicates that *XBP1* expression in CCs is 2x increased compared to CBs. The average for technical duplicates was estimated, normalized on *HPRT* as housekeeping gene, and represented as dark circles; *HPRT* expression (set at 1) is represented by the dotted line; mean and SD are also shown.

**Supplementary Table E1.** Healthy controls.

| Sample | Age | Date of birth | Date of analysis | Exact Age | Sex |
| --- | --- | --- | --- | --- | --- |
| HC212 | ped | 20/10/2003 | 26/10/2015 | 12 | F |
| HC213 | ped | 07/10/2003 | 26/10/2015 | 12 | M |
| HC214 | ped | 20/10/2002 | 26/10/2015 | 13 | M |
| HC215 | ped | 12/09/2002 | 26/10/2015 | 13 | M |
| HC216 | ped | 14/09/2001 | 26/10/2015 | 14 | F |
| HC273 | ped | 22/07/2003 | 15/02/2016 | 12 | F |
| HC275 | ped | 15/05/1999 | 15/02/2016 | 16 | F |
| HC277 | ped | 13/04/2003 | 29/02/2016 | 12 | M |
| HC278 | ped | 15/11/2001 | 29/02/2016 | 14 | F |
| HC280 | ped | 08/07/2002 | 29/02/2016 | 13 | M |
| HC307 | ped | 17/05/2004 | 30/05/2016 | 12 | M |
| HC308 | ped | 08/08/2003 | 30/05/2016 | 12 | M |
| HC309 | ped | 18/08/2002 | 30/05/2016 | 13 | M |
| HC310 | ped | 09/04/2005 | 30/05/2016 | 11 | M |
| HC325 | ped | 09/10/2004 | 14/11/2016 | 12 | F |
| HC326 | ped | 12/02/2004 | 14/11/2016 | 12 | M |
| HC327 | ped | 18/11/2002 | 14/11/2016 | 14 | M |
| HC328 | ped | 09/09/2002 | 14/11/2016 | 14 | F |
| HC331 | ped | 27/01/2005 | 05/12/2016 | 11 | F |
| HC333 | ped | 11/03/2003 | 05/12/2016 | 13 | F |
| HC334 | ped | 06/08/2002 | 05/12/2016 | 14 | F |
| HC335 | ped | 11/09/2000 | 05/12/2016 | 16 | M |
| HC336 | ped | 30/09/2004 | 12/12/2016 | 12 | M |
| HC337 | ped | 28/12/2007 | 12/12/2016 | 8 | M |
| HC344 | ped | 13/04/2004 | 09/01/2017 | 12 | M |
| HC346 | ped | 03/03/2003 | 09/01/2017 | 13 | F |
| HC349 | ped | 13/11/2011 | 25/01/2017 | 5 | M |
| HC350 | ped | 13/08/2008 | 25/01/2017 | 8 | M |
| HC396 | ped | 17/06/2009 | 20/09/2017 | 8 | F |
| HC397 | ped | 27/06/2010 | 21/09/2017 | 7 | F |
| HC398 | ped | 27/07/2013 | 25/09/2017 | 4 | F |
| HC400 | ped | 13/06/2005 | 12/10/2017 | 12 | M |
| HC401 | ped | 28/11/2005 | 12/10/2017 | 11 | M |
| HC431 | ped | 15/11/2005 | 12/04/2018 | 12 | F |
| HC462 | ped | 29/10/2009 | 13/05/2019 | 9 | M |
| HC449 | ped | 01/10/2016 | 22/07/2019 | 2 | F |
| HC409 | ped | 31/12/2004 | 22/11/2017 | 12 | M |
| HC421 | ped | 07/11/2008 | 14/02/2018 | 9 | F |
| HC436 | ped | 01/09/2002 | 16/05/2018 | 15 | F |
| HC359 | ad | 12/05/1990 | 03/05/2017 | 27 | F |
| HC055 | ad | 29/08/1982 | 12/10/2017 | 35 | F |
| HC068 | ad | 12/05/1977 | 12/10/2017 | 40 | F |
| HC145 | ad | 04/08/1981 | 10/05/2017 | 35 | F |
| HC148 | ad | 23/04/1976 | 21/03/2016 | 39 | F |
| HC151 | ad | 03/04/1969 | 08/05/2017 | 46 | F |
| HC159 | ad | 04/11/1990 | 08/05/2017 | 26 | M |
| HC184 | ad | 28/04/1963 | 24/03/2016 | 52 | F |
| HC194 | ad | 24/03/1989 | 05/06/2016 | 27 | F |
| HC198 | ad | 22/11/1996 | 08/10/2015 | 18 | F |
| HC199 | ad | 29/07/1996 | 08/10/2015 | 19 | F |
| HC200 | ad | 22/12/1989 | 08/10/2015 | 25 | F |
| HC211 | ad | 05/03/1992 | 14/10/2015 | 23 | M |
| HC221 | ad | 23/05/1995 | 11/12/2015 | 20 | M |
| HC232 | ad | 10/09/1993 | 18/01/2016 | 22 | M |
| HC233 | ad | 11/08/1996 | 18/01/2016 | 19 | F |
| HC234 | ad | 15/08/1996 | 18/01/2016 | 19 | F |
| HC235 | ad | 18/11/1992 | 18/01/2016 | 23 | M |
| HC236 | ad | 17/10/1995 | 18/01/2016 | 20 | M |
| HC242 | ad | 31/05/1996 | 19/01/2016 | 19 | F |
| HC243 | ad | 05/04/1996 | 19/01/2016 | 19 | F |
| HC247 | ad | 13/02/1996 | 21/01/2016 | 20 | F |
| HC248 | ad | 03/08/1996 | 21/01/2016 | 19 | F |
| HC249 | ad | 13/09/1994 | 25/01/2016 | 21 | M |
| HC250 | ad | 21/06/1995 | 25/01/2016 | 20 | M |
| HC251 | ad | 02/11/1993 | 25/01/2016 | 22 | F |
| HC252 | ad | 17/09/1995 | 25/01/2016 | 20 | F |
| HC253 | ad | 21/03/1996 | 25/01/2016 | 19 | F |
| HC262 | ad | 02/10/1996 | 27/01/2016 | 19 | F |
| HC263 | ad | 25/10/1995 | 27/01/2016 | 20 | F |
| HC268 | ad | 06/10/1996 | 28/01/2016 | 19 | F |
| HC269 | ad | 25/06/1988 | 13/04/2016 | 27 | F |
| HC271 | ad | 26/12/1988 | 09/02/2016 | 27 | F |
| HC276 | ad | 15/03/1980 | 16/02/2016 | 35 | F |
| HC282 | ad | 18/07/1980 | 08/03/2016 | 35 | M |
| HC283 | ad | 01/01/1980 | 08/03/2016 | 36 | M |
| HC284 | ad | 12/03/1994 | 08/03/2016 | 22 | M |
| HC285 | ad | 27/08/1994 | 08/03/2016 | 21 | M |
| HC286 | ad | 25/08/1994 | 08/03/2016 | 21 | F |
| HC301 | ad | 15/03/1980 | 26/04/2016 | 36 | F |
| HC311 | ad | 11/04/1998 | 30/05/2016 | 18 | M |
| HC312 | ad | 10/07/1996 | 08/10/2015 | 19 | M |
| HC319 | ad | 10/12/1987 | 30/08/2016 | 28 | M |
| HC323 | ad | 12/10/1987 | 14/10/2016 | 29 | F |
| HC340 | ad | 06/06/1989 | 14/12/2016 | 27 | M |
| HC351 | ad | 31/12/1993 | 08/08/2018 | 24 | F |
| HC359 | ad | 12/05/1990 | 03/05/2017 | 27 | F |
| HC360 | ad | 21/08/1991 | 15/02/2017 | 25 | M |
| HC368 | ad | 03/12/1984 | 01/03/2017 | 32 | F |
| HC386 | ad | 24/12/1989 | 05/04/2017 | 27 | F |
| HC389 | ad | 23/10/1993 | 11/05/2017 | 23 | M |
| HC390 | ad | 02/11/1983 | 18/05/2017 | 33 | F |
| HC430 | ad | 12/05/1992 | 22/03/2018 | 25 | M |
| HC441 | ad | 03/09/1989 | 08/08/2018 | 28 | F |
| HC442 | ad | 26/11/1968 | 09/08/2018 | 49 | F |
| HC443 | ad | 09/06/1983 | 20/09/2018 | 35 | F |
| HC448 | ad | 19/07/1972 | 12/12/2018 | 46 | F |
| HC456 | ad | 08/02/1994 | 04/04/2019 | 25 | F |
| HC457 | ad | 06/12/1991 | 19/04/2019 | 27 | M |
| HC458 | ad | 13/10/1995 | 23/04/2019 | 23 | F |
| HC463 | ad | 11/07/1991 | 07/06/2019 | 28 | F |
| HC464 | ad | 04/07/1991 | 18/06/2019 | 28 | F |
| HC458 | ad | 13/10/1995 | 21/06/2019 | 23 | F |
| HC351 | ad | 31/12/1993 | 25/06/2019 | 26 | F |
| HC440 | ad | 05/11/1991 | 27/06/2019 | 27 | F |
| HC475 | ad | 29/10/1992 | 12/11/2019 | 27 | F |

**Supplementary Table E2.**
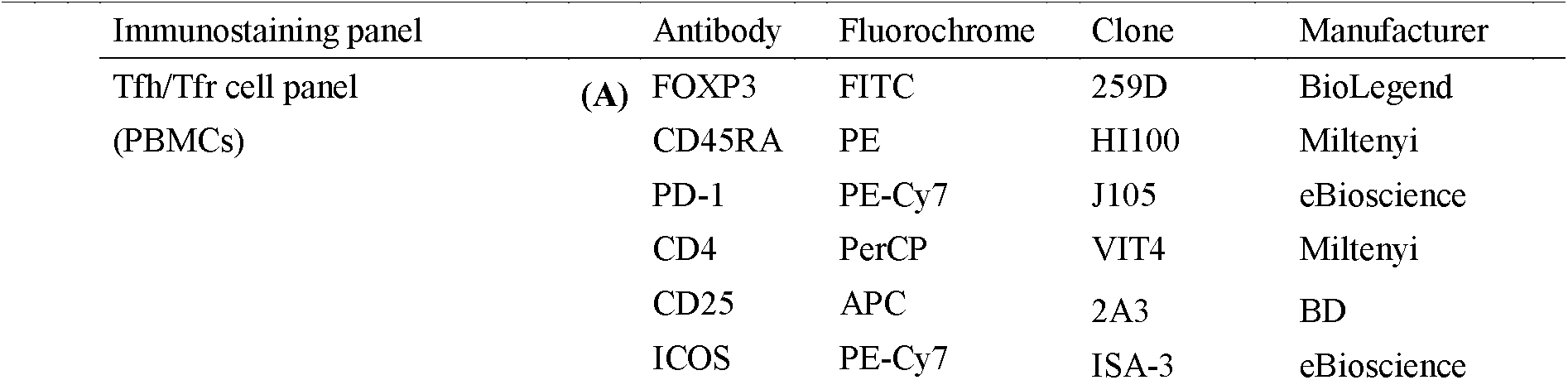
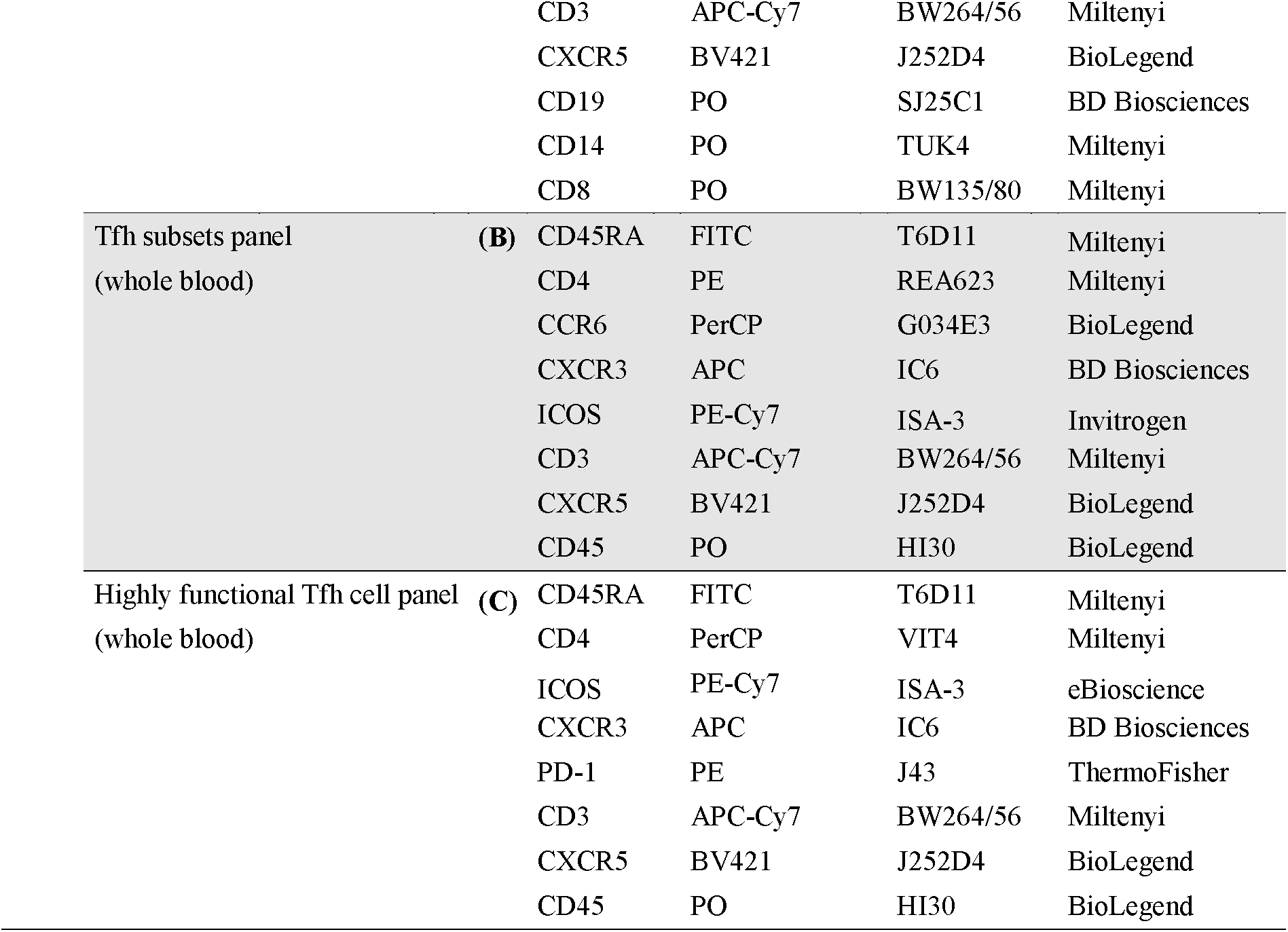
Antibodies and immunostaining panels used for whole blood and PBMC.

**Supplementary Table E3.** Antibodies and immunostaining panels used for PBMC and spleen.

| Immunostaining panel |  | Antibody | Fluorochrome | Clone | Manufacturer |
| --- | --- | --- | --- | --- | --- |
| B cells<br>(Mix 1) | **(A)** | IgM | FITC | G20-127 | BD Biosciences |
|  |  | CD21 | PE | B-LY4 | BD Biosciences |
|  |  | CD27 | APC | M-T271 | BD Biosciences |
|  |  | CD38 | PerCP-Cy5.5 | HIT2 | BD Biosciences |
|  |  | CD19 | PE-Cy7 | SJ 25C1 | BD Biosciences |
|  |  | CD24 | PB | SN3 | EXBIO |
|  |  | IgD | BIO | IA6-2 | BD Biosciences |
|  |  | Streptavidin | PO | - | ThermoFisher |
| B cells<br>(Mix 2) | **(B)** | IgA | FITC | polyclonal | Jackson ImmunoResearch |
|  |  | CD21 | PE | B-LY4 | BD Biosciences |
|  |  | CD27 | APC | M-T271 | BD Biosciences |
|  |  | CD38 | PerCP-Cy5.5 | HIT2 | BD Biosciences |
|  |  | CD19 | PE-Cy7 | SJ 25C1 | BD Biosciences |
|  |  | CD24 | PB | SN3 | EXBIO |
|  |  | IgD | BIO | IA6-2 | BD Biosciences |

|  |  | Streptavidin | PO | - | ThermoFisher |
| --- | --- | --- | --- | --- | --- |
| B cells<br>(Mix 3) | (C) | IgG | FITC | polyclonal | Jackson ImmunoResearch |
|  |  | CD21 | PE | B-LY4 | BD Biosciences |
|  |  | CD27 | APC | M-T271 | BD Biosciences |
|  |  | CD38 | PerCP-Cy5.5 | HIT2 | BD Biosciences |
|  |  | CD19 | PE-Cy7 | SJ 25C1 | BD Biosciences |
|  |  | CD24 | PB | SN3 | EXBIO |
|  |  | IgD | BIO | IA6-2 | BD Biosciences |
|  |  | Streptavidin | PO | - | ThermoFisher |
| Tfh cells | (D) | Ki67 | FITC | B56 | BD Biosciences |
|  |  | CD57 | PE | TB03 | Miltenyi |
|  |  | BCL-6 | APC | 7D1 | BioLegend |
|  |  | CD4 | PerCP-Cy5.5 | VIT4 | Miltenyi |
|  |  | PD-1 | PE-Cy7 | J105 | eBiosciences |
|  |  | CD3 | APC-Cy7 | BW264/56 | Miltenyi |
|  |  | CXCR5 | BV421 | J252D4 | BioLegend |
|  |  | CD8 | PO | BW135/80 | Miltenyi |
| Tfh, centroblasts,<br>centrocytes | (E) | CD3 | APC-Cy7 | BW264/56 | Miltenyi |
|  |  | CD4 | PerCP | VIT4 | Miltenyi |
|  |  | CXCR5 | BV421 | J252D4 | BioLegend |
|  |  | PD-1 | PE | J43 | ThermoFisher |
|  |  | CD27 | APC | M-T271 | BD Biosciences |
|  |  | CD38 | PerCP-Cy5.5 | HIT2 | BD Biosciences |
|  |  | CXCR4 | PE-Cy7 | 12G5 | Miltenyi |

**Supplementary Table E4.**
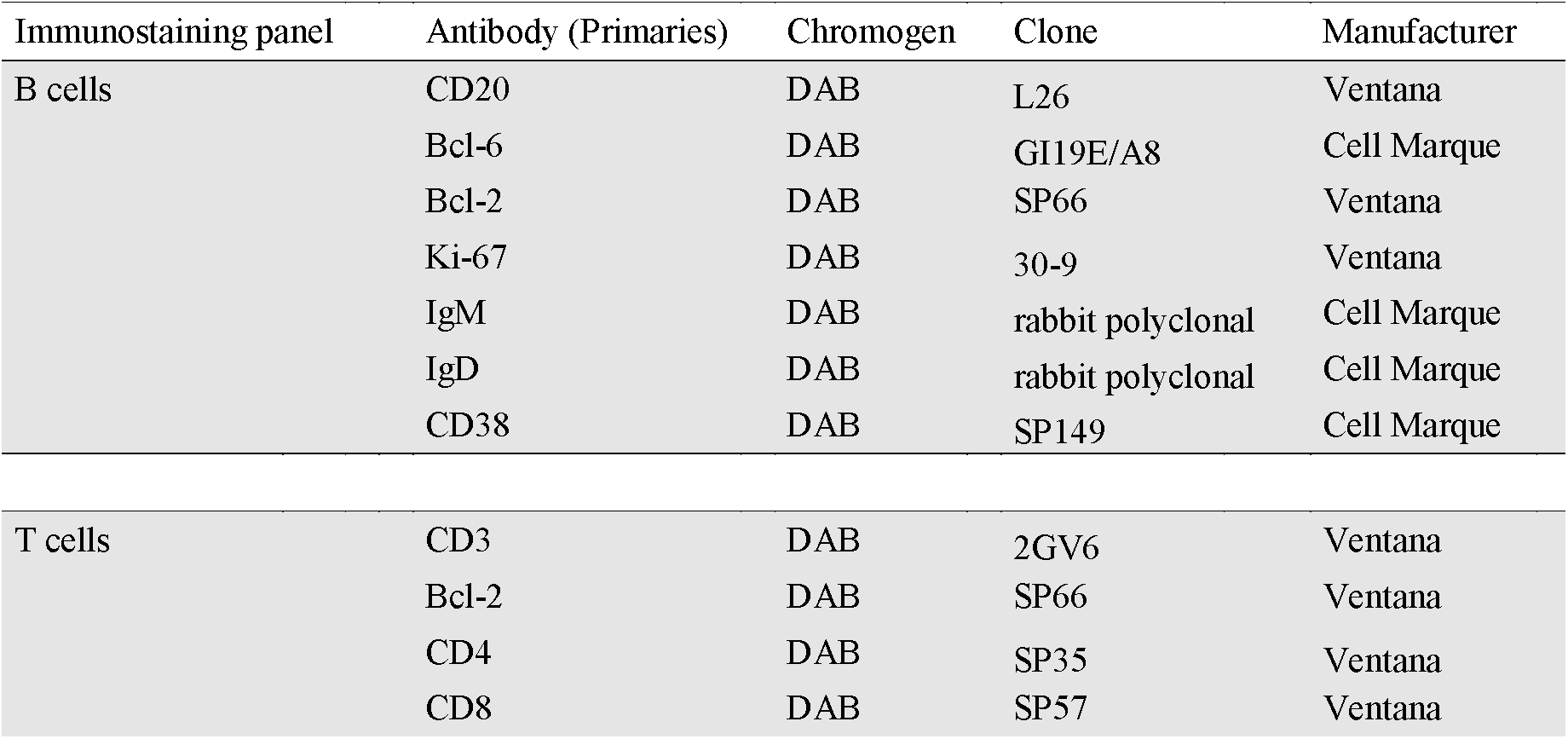
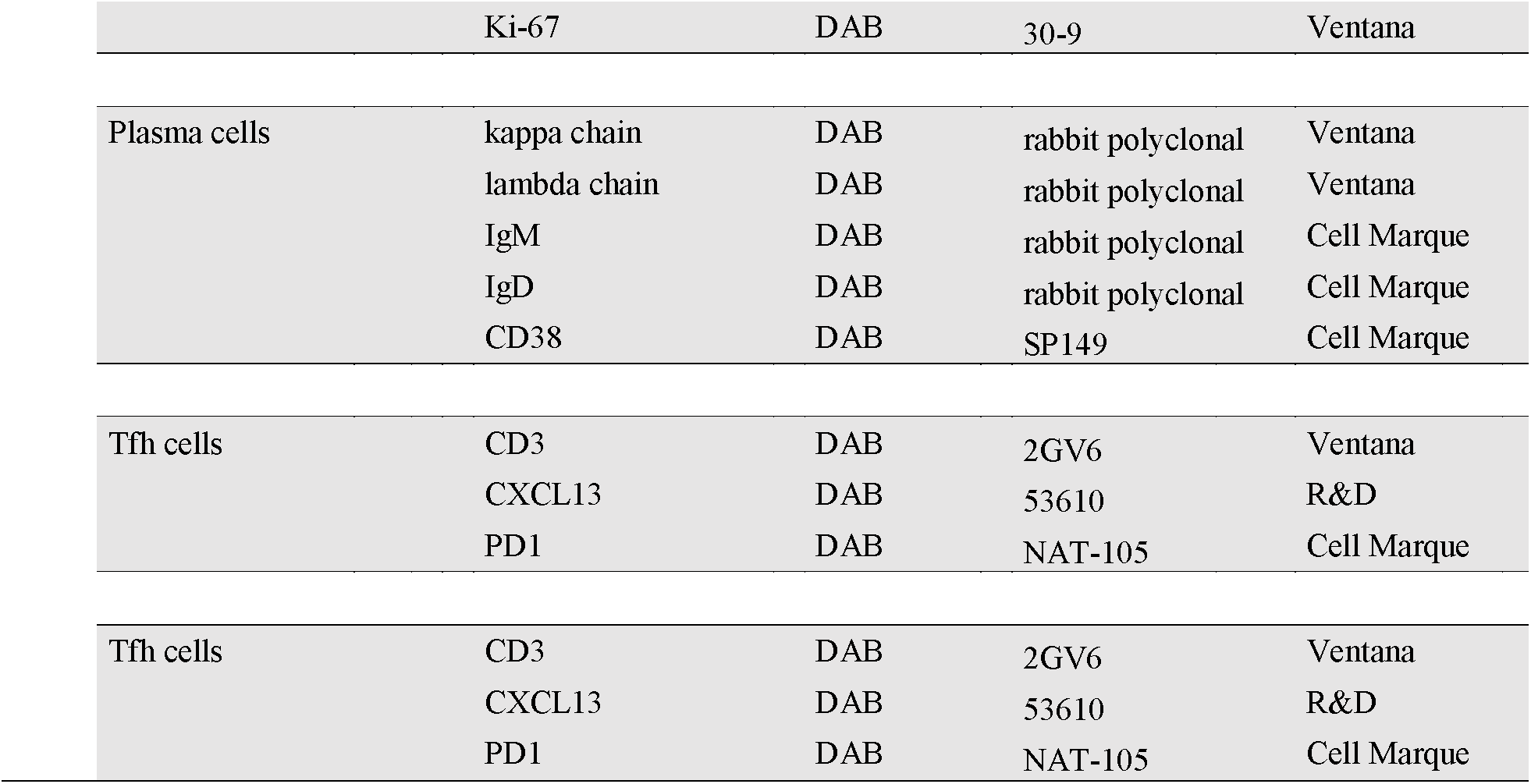
Antibodies and immunostaining panel used for immunohistochemistry.

**Supplementary Table E5.**
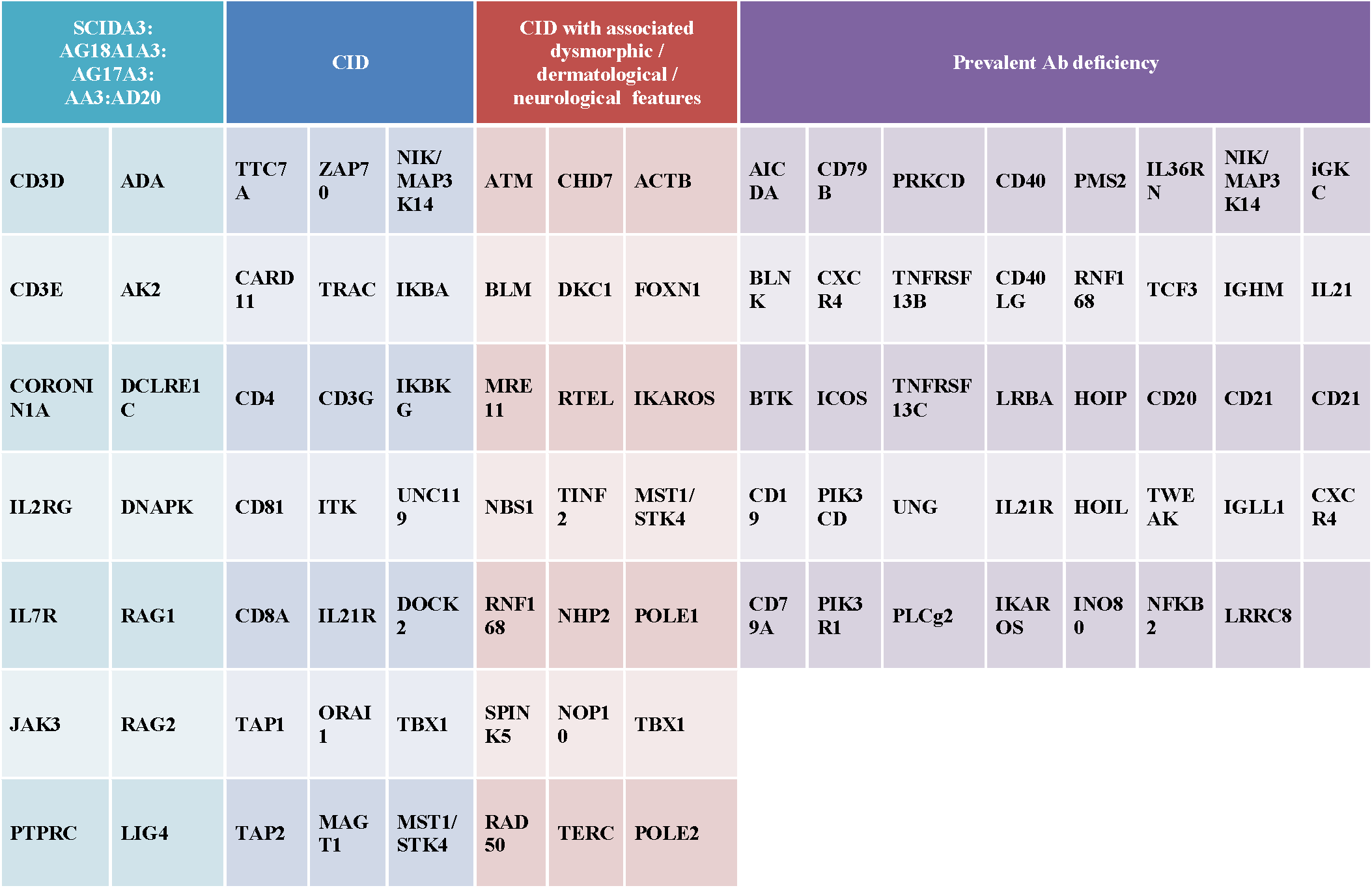
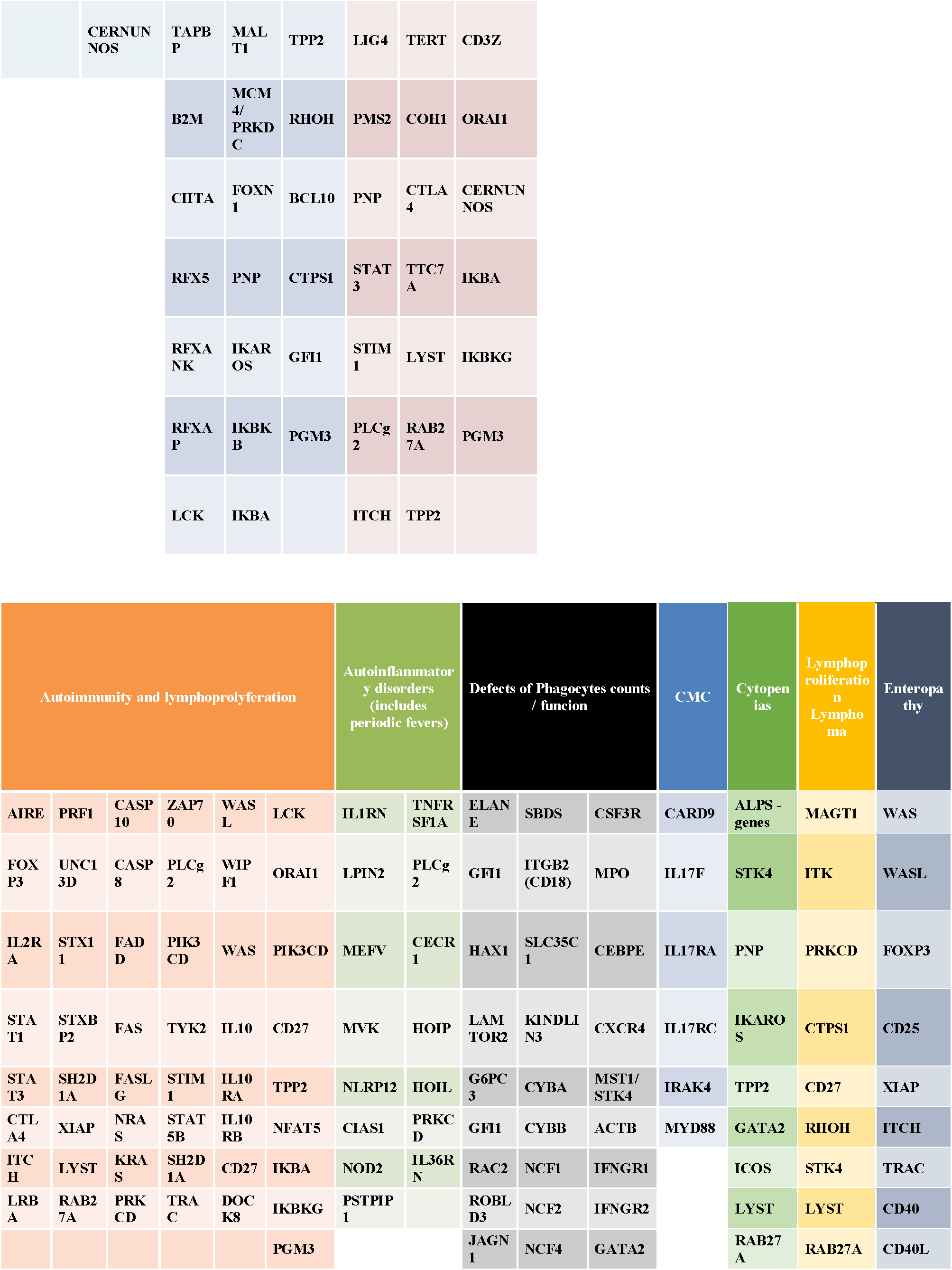
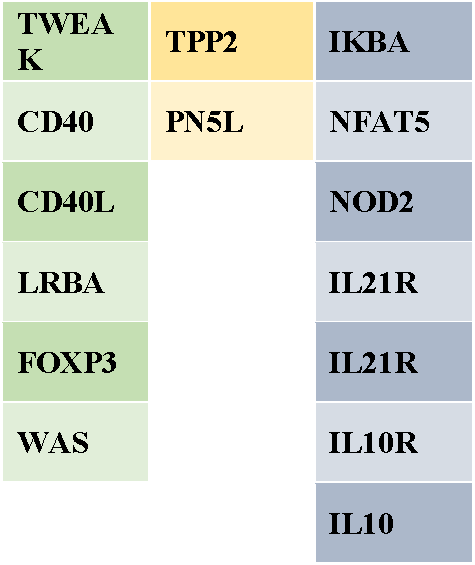
Gene pipeline for the discovery of causative mutations.

**Supplementary Table E6.**
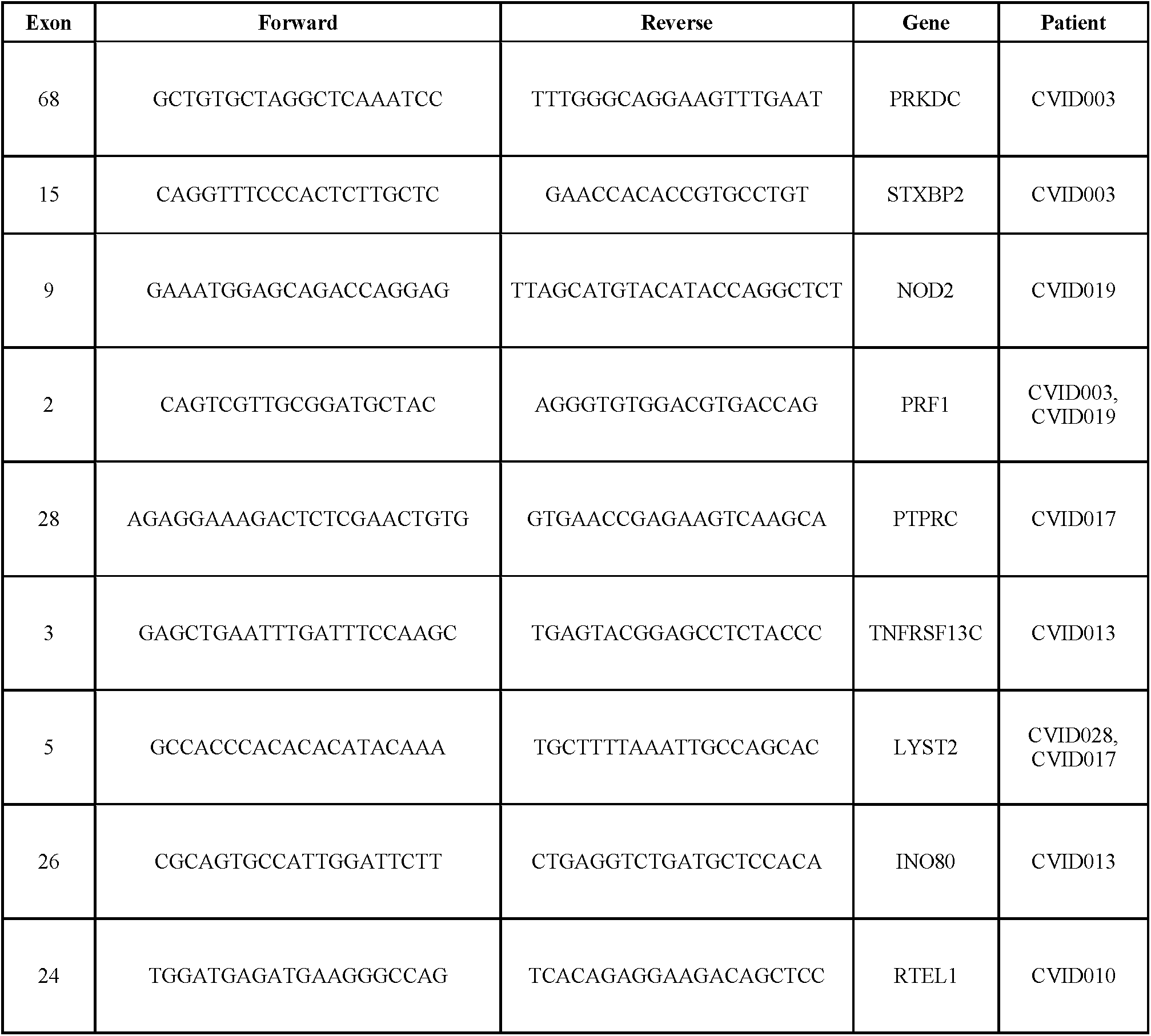
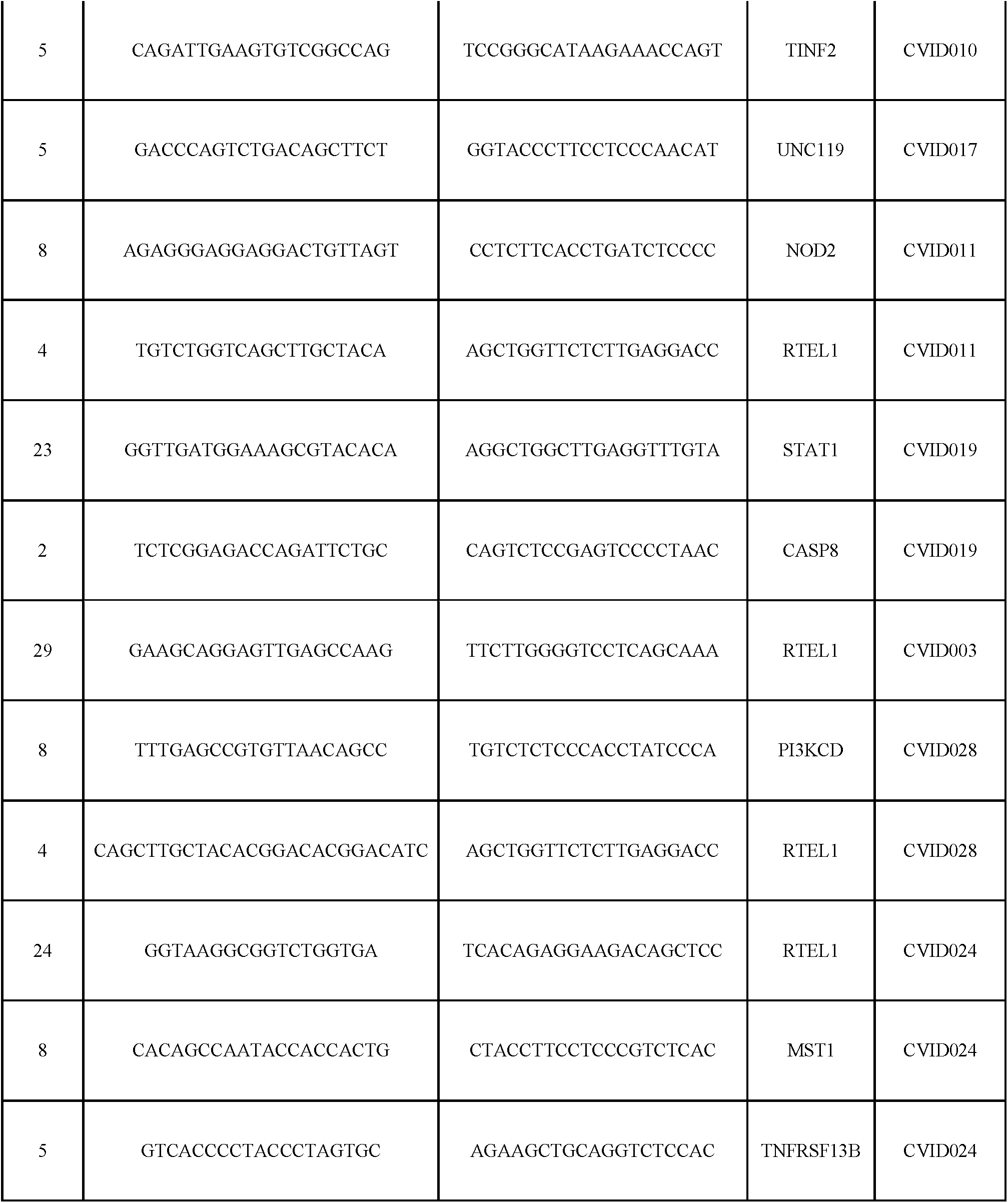
Primers used for amplification and sequencing of genomic DNA.

**Supplementary Table E7.**
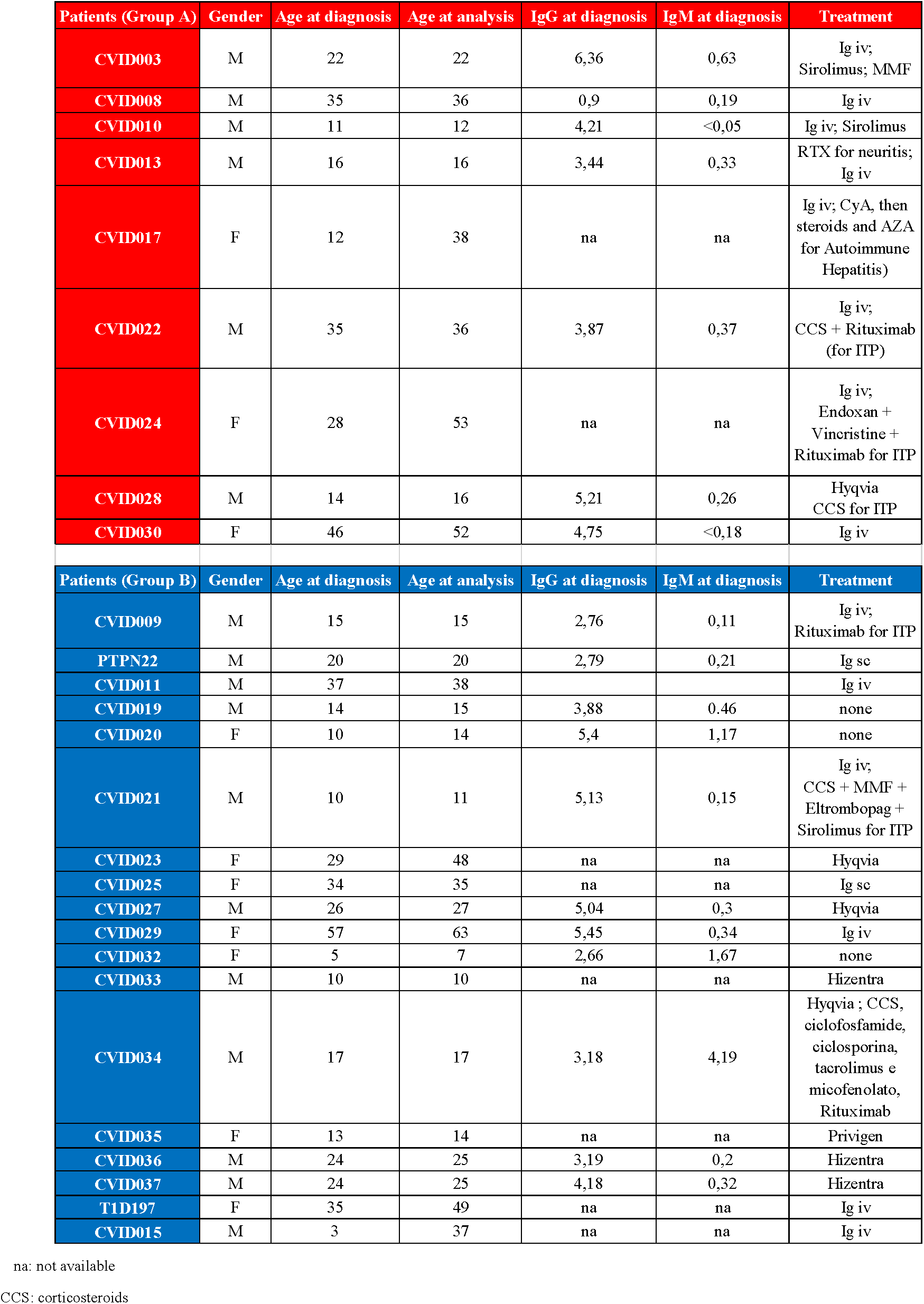
Therapeutic and clinical features of CVID patients of group A and B.

